# Synaptotagmin-7 places dense-core vesicles at the cell membrane to promote Munc13-2- and Ca^2+^-dependent priming

**DOI:** 10.1101/2020.11.02.365478

**Authors:** Bassam Tawfik, Joana S. Martins, Sébastien Houy, Cordelia Imig, Paulo S. Pinheiro, Sonja M. Wojcik, Nils Brose, Benjamin H. Cooper, Jakob B. Sørensen

**Author notes:** Equal contribution (Joana S. Martins, Sébastien Houy, Cordelia Imig). Correspondence to: Jakob B. Sørensen Department of Neuroscience University of Copenhagen Blegdamsvej 3B 2200 Copenhagen N Denmark.

## Abstract

The functional consequences of the co-expression of synaptotagmin-1 and synaptotagmin-7 are unclear. We show that when present separately, synaptotagmin-1 and synaptotagmin-7 act as standalone fast and slow Ca^2+^-sensors for vesicle fusion in mouse chromaffin cells. When present together, synaptotagmin-7 stimulates Ca^2+^-dependent vesicle priming and inhibits depriming. The priming effect of Synaptotagmin-7 extends to the Readily Releasable Pool, whose fusion is executed by synaptotagmin-1, indicating synergistic action of the two Ca^2+^-sensors, although they are only partially colocalized. Synaptotagmin-7 promotes ubMunc13-2-dependent priming and the absence of synaptotagmin-7 renders phorbolesters less effective in stimulating priming, although synaptotagmin-7 independent priming is also observed. Morphologically, synaptotagmin-7 places vesicles in close membrane apposition (< 6 nm); in its absence vesicles accumulate out of reach of the fusion complex (20-40 nm). We suggest that a synaptotagmin-7-dependent movement toward the membrane is involved in Munc13-2/phorbolester/Ca^2+^-dependent priming and sets the stage for fast and slow exocytosis triggering.

## Introduction

Neurotransmitter or hormone release requires a basal membrane fusion machinery, and one or more Ca^2+^-sensors to link fusion to the electrical activity of the cell. The machinery driving vesicle-to-membrane fusion consists of the SNAREs (Fang and Lindau, 2014; Jahn and Fasshauer, 2012), associated proteins Munc18 and Munc13 necessary for SNARE-complex assembly (Rizo and Xu, 2015), and Ca^2+^-sensors of the synaptotagmin (Syt) family (Pinheiro et al., 2016) together with complexins (Makke et al., 2018; Trimbuch and Rosenmund, 2016). Syts harbor two C2-domains, which can bind to Ca^2+^ and phospholipids (in 8 of the 17 Syt isoforms present in the mammalian genome), and to SNAREs (Sudhof, 2002). Analysis in expressing cells showed that Syt-1, Syt-2, and Syt-9 trigger fast, synchronous release (Xu et al., 2007). Synaptotagmin-7 (Syt-7) displays the slowest membrane binding/unbinding kinetics of all the synaptotagmins (Hui et al., 2005), and the highest Ca^2+^-affinity during lipid binding (Bhalla et al., 2005; MacDougall et al., 2018; Sugita et al., 2002), although the apparent Ca^2+^-affinities vary with the experimental system and lipid composition (in terms of half-maximal lipid-binding, Syt-7: 0.3-2 μM; Syt-1: 10-150 μM (Bhalla et al., 2005; Sugita et al., 2002)). Syt-7 is expressed in high amounts in the brain (Sugita et al., 2001) and the adrenal medulla (Schonn et al., 2008) where it is typically found co-expressed with Syt-1. The co-existence of two Syts with very different kinetics and Ca^2+^-affinities in the same cell raises the question whether they act autonomously/additive, compete with each other, or cooperate in a shared mechanism (Walter et al., 2011).

When Syt-1 is deleted, fast release is eliminated and residual slow/asynchronous release is driven by Syt-7 (Bacaj et al., 2013; Schonn et al., 2008), which is consistent with an additive function of the two sensors. However, the function of Syt-7 in the presence of the faster Syt isoform (Syt-1 or Syt-2) is not clear. In the presence of Syt-1 or Syt-2, Syt-7 does not affect synchronous or asynchronous release following a single action potential in cultured neurons (Liu et al., 2014; Weber et al., 2014), in the Calyx of Held synapse (Luo and Sudhof, 2017) or in cerebellar basket cell synapses (Chen et al., 2017). However, during high frequency stimulation Syt-7 adds a sustained or tonic component of release in a number of different depressing synapses (Bacaj et al., 2013; Chen et al., 2017; Liu et al., 2014; Luo and Sudhof, 2017; Turecek et al., 2017; Wen et al., 2010). Reduced sustained and fast release was first described in the Syt-7 knock-out (KO) chromaffin cells (Schonn et al., 2008). Liu et al. suggested that Syt-7 acts as an upstream Ca^2+^-sensor that speeds up vesicle recruitment during train stimulation, resulting in sustained release during trains while the Ca^2+^ concentration is high (Liu et al., 2014). The function in priming was supported by delayed calcium-dependent recovery after high frequency or sucrose stimulations in cultured glutamatergic hippocampal neurons from the Syt-7 KO (Liu et al., 2014). Conversely, Bacaj and collaborators did not find differences in recovery in Syt-7 KO hippocampal neurons; instead, they reported that Syt-7 acts together with Syt-1 to ensure full capacity of the primed vesicle pool, which implies a function in stabilizing primed vesicles (Bacaj et al., 2015). Syt-7 also supports asynchronous glutamate release from principal cells onto Martinotti cells (Deng et al., 2020) and Syt-7 is necessary for short-term synaptic facilitation (Jackman et al., 2016). After prolonged stimulation trains, or in the presence of manipulations exacerbating asynchronous release, Syt-7 directs vesicles towards a slow endocytosis pathway (Li et al., 2017; Virmani et al., 2003). Thus, Syt-7 plays multiple roles in exocytosis and endocytosis (Chen and Jonas, 2017; Huson and Regehr, 2020; MacDougall et al., 2018).

Chromaffin cells were the first cells in which vesicle priming was shown to be calcium-dependent (Bittner and Holz, 1992; von Ruden and Neher, 1993), which was later found to apply also to neurons (Dittman and Regehr, 1998; Gomis et al., 1999; Schneggenburger et al., 2002; Stevens and Wesseling, 1998; Wang and Kaczmarek, 1998). Chromaffin cells also display both fast and slow release phases when stimulated by a strong stimulus, such as Ca^2+^ uncaging (Heinemann et al., 1993). Here, we used adrenal chromaffin cells, mouse knockouts, electrophysiology, and high-resolution 3D electron tomography, to investigate whether in this cellular system the two Ca^2+^-sensors Syt-1 and Syt-7 can be said to act independently, or whether they are interdependent - i.e. engaging in either cooperative or competitive interplay (Walter et al., 2011).

## Results

Chromaffin cells offer distinct advantages in the study of neurosecretion (Neher, 2018). Since chromaffin cells do not have a limited number of release sites, Ca^2+^-dependent priming leads to a large and readily measurable increase in the size of the pool of primed large dense-core vesicles (LDCV) when pre-stimulation [Ca^2+^] exceeds 100-200 nM (Voets, 2000). When combined with intracellular Ca^2+^-control via Ca^2+^ -uncaging, this makes it possible to accurately titrate the Ca^2+^-dependence of priming in the steady state (Voets, 2000), something which has not been achieved in other cell types. Here, we define ‘priming’ as the reaction (re-)filling the releasable vesicle pools; a clear kinetic distinction between priming and fusion when triggered by a common stimulus (Ca^2+^) requires that fusion is >5-10-fold faster than priming; this requirement is fulfilled using Ca^2+^-uncaging, which increases the fusion rate abruptly. For these reasons, we used chromaffin cells and Ca^2+^-uncaging to investigate the involvement of Syt-1 and Syt-7 in the Ca^2+^-dependence of priming and fusion triggering.

### When present alone, Syt-1 and Syt-7 act as kinetically distinct fusion triggers

We first studied the function of each Syt isoform in isolation, by expressing Syt-1 and Syt-7 individually using lentiviral vectors (see Materials and Methods) in chromaffin cells cultured from Syt-1/Syt-7 double KO (DKO) mouse embryos (Schonn et al., 2008). We stimulated secretion using Ca^2+^-uncaging to abruptly raise the intracellular Ca^2+^ concentration ([Ca^2+^]_i_) from the sub-μM range to around 20-30 μM (Fig. 1A). This mode of stimulation causes LDCV fusion at a spatially homogeneous [Ca^2+^]_i_, and allows the distinction between fusion kinetics and vesicle pool size. Of these, vesicle pool sizes are affected by priming reactions, which take place before arrival of the fusion trigger (Ca^2+^), whereas fusion kinetics is determined by events downstream of Ca^2+^ arrival. Exocytosis was monitored simultaneously by membrane capacitance measurements, which assesses the plasma membrane area, and amperometry, which measures oxidizable neurotransmitters (mainly adrenaline). Upon Ca^2+^ uncaging, DKO cells displayed a very small response (Fig. 1A) (Schonn et al., 2008). In contrast, when either Syt-1 or Syt-7 was expressed, robust secretion resulted (Fig. 1A). However, the kinetics of secretion were very different: Syt-1 supported a small rapid burst of secretion, with a fusion time constant around 10-20 ms (Fig. 1A-C). In contrast, Syt-7 stimulated a larger burst of secretion, but with a much slower fusion time constant, around 500 ms (Fig. 1A-C). These two components correspond kinetically to the previously identified Readily Releasable Pool (RRP) and Slowly Releasable Pool (SRP) (Voets, 2000). Fitting individual responses with a sum of two exponentials (representing the RRP and the SRP) and a straight line (for the sustained component) allowed us to identify the size and fusion kinetics of both the RRP and SRP (Fig. 1B-C; see Materials and Methods). Syt-1 supported fast secretion (i.e. a RRP, fusion time constant 13.4 ± 1.47 ms), but not slow-burst secretion (i.e. a SRP), whereas for Syt-7 it was opposite (SRP fusion time constant: 500 ± 48.9 ms; Fig. 1B-C). The data also show that the slow burst became faster in the presence of Syt-1, and the fast burst was slower in the presence of Syt-7 (Fig. 1C); thus, the effect of the Syts extend to both phases of release. The Syt-1/Syt-7 DKO apparently supported a very small RRP with fast time constant (Fig. 1B-C); however, caution should be exercised when considering the kinetics of a pool this small (3.4 ± 0.9 fF, corresponding to ~3.5 vesicles (Pinheiro et al., 2014)). Both isoforms supported a near-linear sustained release component (Fig. 1 A,D). Fig. 1A (bottom panel) shows amperometric current and charge; note that the charge is consistent with the capacitance measurements, showing faster kinetics for Syt-1 driven release (quantification of total integrated amperometry is found in Fig. S1A). Thus, expression of either Syt-1 or Syt-7 can re-establish release in Syt-1/7 DKO cells and Syt-1 triggers faster fusion than Syt-7.

**Figure 1.**
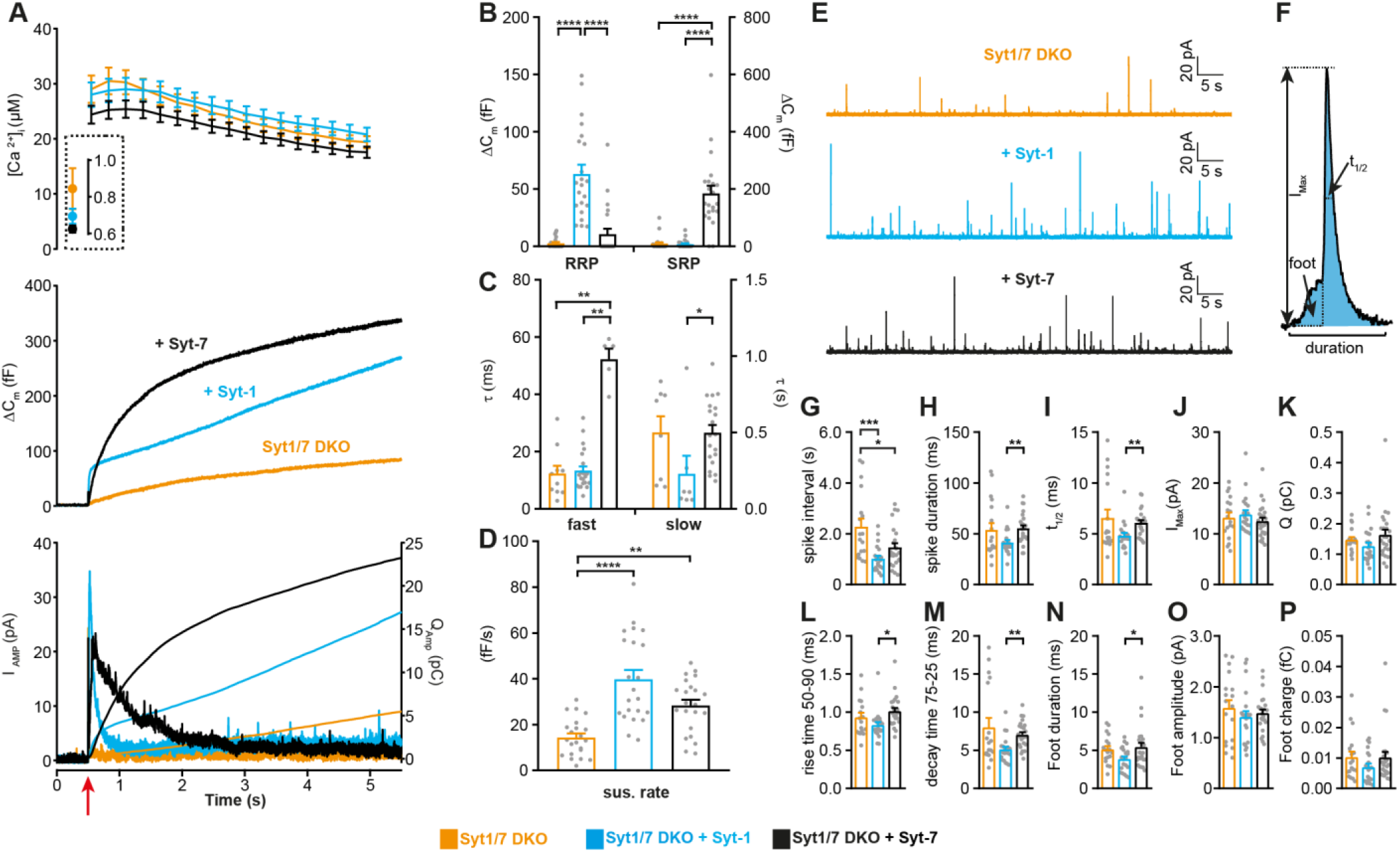
Syt-1 and Syt-7 are stand-alone calcium-sensors with different kinetics. **A** Calcium uncaging experiment in Syt-1/Syt-7 DKO cells (orange) and in DKO cells overexpressing Syt-7 (black) or Syt-1 (blue). Top panel: [Ca^2+^] before (insert) and after calcium uncaging (uncaging flash at red arrow, bottom panel). Middle panel: capacitance traces (mean of all cells) show that the secretion is potentiated more (higher amplitude) by Syt-7 expression, but the kinetics of the secretory burst is faster after Syt-1 expression. Bottom panel: Mean amperometry (left ordinate axis) and mean integrated amperometry (right ordinate axis). Note that the integrated amperometric traces agree very well with the capacitance traces. **B** Sizes of the RRP and SRP. **C** Time constants, τ, of fusion for fast (i.e. RRP) and slow (i.e. SRP) secretion. **D** Sustained rates of secretion. Data information: In (A-D), data with error bars are presented as mean ± SEM; in (A), the traces are the mean of all cells. *: p<0.05; **: p<0.01; ***: p<0.001; ****: p<0.0001. Kruskal-Wallis test with Dunn’s post-hoc test. Number of cells, DKO: N = 23 cells; DKO + Syt-1: N = 22 cells; DKO + Syt-7: N = 21 cells. **E** Amperometric currents induced by infusion of ~5 μM Ca^2+^ into the cell via a patch pipette. Syt-1/Syt-7 DKO cells, DKO cells expressing either Syt-1 (blue trace) or Syt-7 (black trace). **F** Single amperometric spike, indicating measurement of peak current (I_max_), total charge (Q, by integration), duration at half maximum (t_1/2_), and total duration of spike. The foot signal, which reports on the fusion pore before it expands, is indicated. **G** The spike interval **H** Spike duration. **I** Duration at half maximum (t_1/2_). **J** Peak current (I_max_). **K** Total charge of foot and spike (Q). **L** Spike 50-90% rise time. **M** Spike 75-25% decay time. **N** Duration of foot signal. **O** Amplitude of foot signal. **P** Charge of foot signal. The spike interval was significantly decreased by expression of either Syt-1 or Syt-7 in DKO cells. The shape parameters show that spikes have faster dynamics in the presence of Syt-1 than with Syt-7. Data information: In (G-P), data are presented as mean ± SEM. *: p<0.05; **: p<0.01; *** p<0.001. In (G, L, N): One-way ANOVA with post-hoc Tukey’s test. In (H, I, M): Kruskal-Wallis test with post-hoc Dunn's test. The spike interval (G) and the duration of foot signal (N) were log-transformed before statistical testing. Number of cells: DKO: N = 18 cells DKO + Syt-1: N = 21 cells; DKO + Syt-7: N = 24 cells.

If the Syts act as stand-alone calcium-sensors, they might be expected to differently affect single-vesicle fusion events. This can be investigated by single-vesicle amperometry, where each fusion event is detected as a spike in the oxidation current (Bruns, 2004). However, previous investigations in mouse chromaffin cells failed to find differences between Syt-1 KO and WT (Voets et al., 2001) or Syt-7 KO and WT spike shapes (Segovia et al., 2010). However, a difference between Syt-1 and Syt-7 might be detectable when comparing DKO cells overexpressing Syt-1 and Syt-7 side-by-side. To elicit sustained secretion at an intermediate frequency – so that single spikes can be resolved – we infused overexpressing DKO cells with an intermediate Ca^2+^ concentration (~5 μM) via the patch pipette. A semi-automatic detection algorithm was used to identify spikes and quantify their shape (Mosharov, 2008). Expression of either Syt-1 or Syt-7 increased the spike frequency – or, equivalently, reduced the spike interval (Fig. 1E, G). This increase in frequency was consistent with the increase in sustained released in uncaging experiments (Fig. 1D); indeed, Ca^2+^-infusion induces sustained release, but does not allow the build-up and abrupt fusion of standing pools of primed LDCVs. When analyzing the shape of the amperometric spikes, Syt-1 caused a shorter spike duration than Syt-7 (Fig. 1H), a faster half-life (Fig. 1I), a faster rise time (Fig. 1L), a faster decay time (Fig. 1M), and a shorter duration of the so-called ‘foot’ signal (Fig. 1N). The foot signal measures the duration of the fusion pore prior to pore expansion (Chow et al., 1992). A longer foot signal following overexpression of Syt-7 compared to Syt-1 was previously found in PC12-cells (Zhang et al., 2010). These findings agree with and expand on data obtained using optical means (TIRF-microscopy), which showed a longer duration of the exocytotic event driven by Syt-7 (Bendahmane et al., 2020; Rao et al., 2014; Rao et al., 2017). Overall, these data establish the status of Syt-1 and Syt-7 as stand-alone fusion sensors and demonstrates that Syt-1 supports faster fusion than Syt-7 on both the population and the single-vesicle level.

### Syt-7 promotes Ca^2+^-dependent priming of slow and fast release (SRP and RRP)

To study the impact of Syt-7 on release in the presence of Syt-1, we used Syt-7 KO chromaffin cells where endogenous levels of Syt-1 remained unchanged (see below: Fig. 9B, Fig. S9B, E). We compared Syt-7 KO cells to WT cells, and Syt-7 KO cells overexpressing Syt-7 (i.e. rescue experiments) using lentiviral transduction. If there is interaction between Syt-7 and Syt-1, it likely involves the different Ca^2+^-affinities: Syt-7 binds to Ca^2+^ with half-maximal binding around or below 1 μM (0.3-2 μM), whereas Syt-1 binds Ca^2+^ at higher concentrations (10-150 μM) (Bhalla et al., 2005; Sugita et al., 2002). Interaction between the isoforms should be revealed when increasing the [Ca^2+^]_i_ in two steps: first, an increase from resting levels to an intermediate value (below 1 μM); second, a rapid increase to higher values (>15 μM). This should allow a sizeable fraction of the Syt-7 molecules to bind Ca^2+^ at the intermediate step, whereas Syt-1 would not bind until after the second stimulus. This can be achieved by accurate adjustment of the [Ca^2+^]_i_ using caged-Ca^2+^ (nitrophenyl-EGTA) infused through a patch-pipette, and a monochromator Xenon lamp oscillating between 350 and 380 nm, which slowly uncages Ca^2+^, thereby raising the pre-stimulation [Ca^2+^]_i_ while [Ca^2+^]_i_ is simultaneously measured by fura dyes (Voets, 2000), to allow on-line control. After spending ~20 s at the intermediate calcium concentration, a rapid uncaging stimulus was delivered by a UV flash lamp; the resulting secretion constitutes the output of the experiment and is shown in Figure 2.

**Figure 2.**
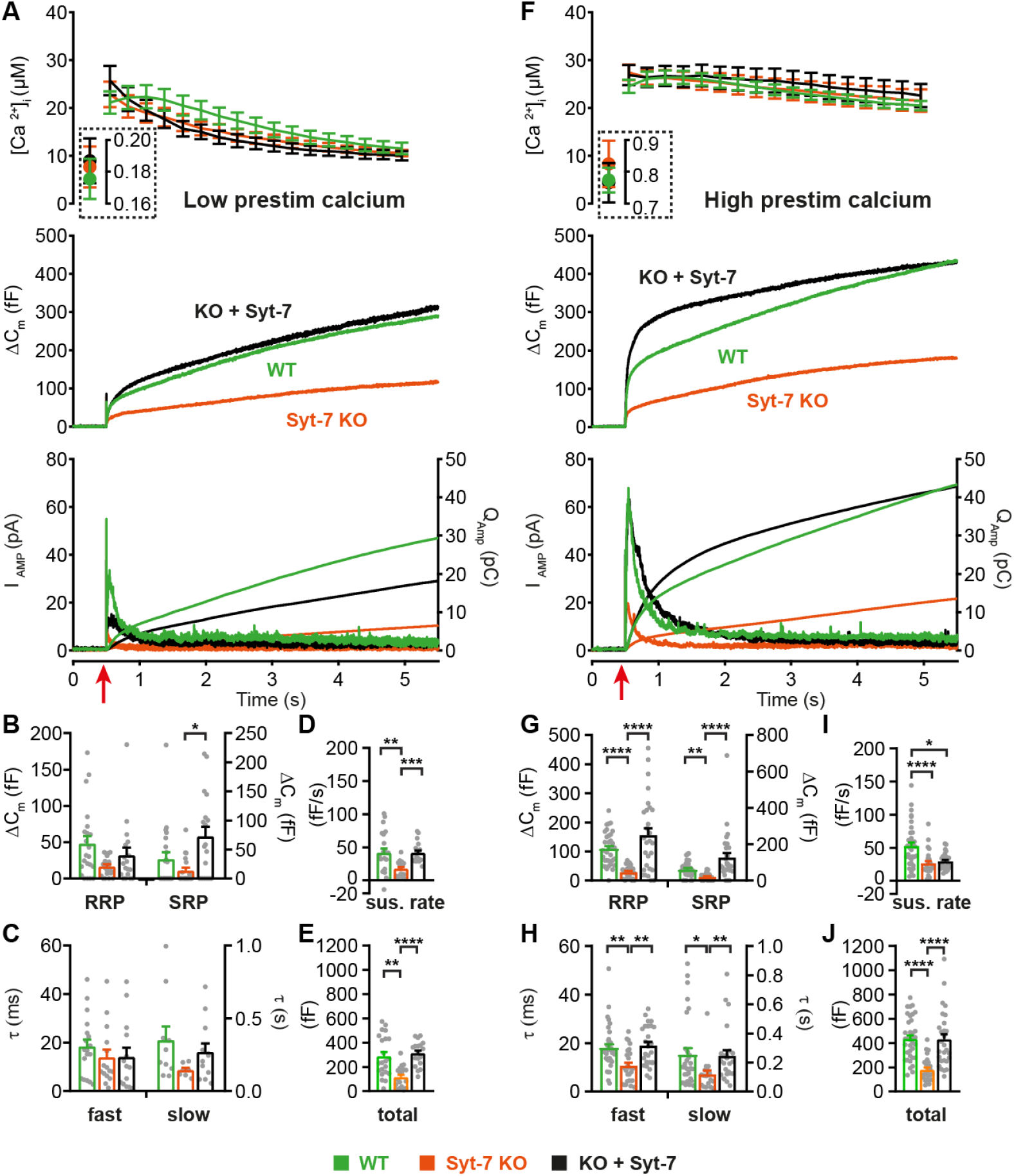
Syt-7 potentiates primed vesicle pool sizes at higher prestimulation [Ca^2+^]. **A** Calcium uncaging experiment from low prestimulation [Ca^2+^] in WT cells (green), Syt-7 KO cells (vermilion) and in Syt-7 KO cells overexpressing Syt-7 (black traces). Panels are arranged as in Fig. 1A. **B** Sizes of the RRP and SRP. **C** Time constants of fusion for fast (i.e. RRP) and slow (i.e. SRP) secretion. **D** Sustained rates of secretion. **E** Total capacitance increase. **F** Calcium uncaging experiment from high prestimulation [Ca^2+^] in WT cells (green), Syt-7 KO cells (vermilion) and in Syt-7 KO cells overexpressing Syt-7 (black traces). Panels arranged as in Fig. 1A. **G** Sizes of the RRP and SRP. **H** Time constants of fusion for fast (i.e. RRP) and slow (i.e. SRP) secretion. **I** Sustained rates of secretion. **J** Total capacitance increase. When stimulated from high prestimulation [Ca^2+^], Syt-7 expression potentiated RRP and SRP size. Data information: In (A-J) data with error bars are presented as mean ± SEM; in (A and F), the traces are the mean of all cells. * p<0.05; ** p<0.01; *** p<0.001; **** p<0.0001. Kruskal-Wallis test with Dunn’s post-hoc test. Number of cells in (A-E): Syt-7 WT: N = 22 cells; Syt-7 KO: N = 19 cells; Syt-7 KO + Syt-7: N = 18 cells, in (F-J) Syt-7 WT: N = 36 cells; Syt-7 KO: N = 27 cells; Syt-7 KO + Syt-7: N = 28 cells. Note that in cases where a cell did not have a given pool (SRP or RRP), the size of that pool was set to zero, and no time constant was estimated.

We first performed uncaging experiments from a relatively low intermediate (‘pre-stimulation’) [Ca^2+^] (Fig. 2A-E) of around 180 nM. Under these conditions, knockout of Syt-7 caused a decrease in secretion as measured by both capacitance and amperometry (Fig. 2A, E; Fig. S1B), whereas re-expression of Syt-7 caused rescue of secretion to WT levels (Fig. 2A, E). Kinetic analysis identified (statistically non-significant) reductions in the average RRP and SRP size in the absence of Syt-7, whereas rescue by Syt-7 overexpression generated an even larger SRP than that observed for endogenous Syt-7 levels (Fig. 2B). Syt-7 expression did not significantly affect the kinetics of slow or fast release (Fig. 2C). The sustained component of release was significantly reduced by the absence of Syt-7 and rescued upon re-expression (Fig. 2D). Expression analysis by immunofluorescence showed that lentiviral expression of the WT Syt-7 results in levels ~2-fold higher than endogenous Syt-7 (Fig. S2A-B).

Raising pre-stimulation [Ca^2+^] to ~0.8 μM resulted in markedly more secretion in Syt-7 expressing cells (Fig. 2F, J), consistent with Ca^2+^-dependent vesicle priming. Under these conditions, both fast (RRP) and slow (SRP) burst secretion were strongly and significantly decreased by Syt-7 knockout and rescued by Syt-7 expression (Fig. 2G, Fig. S1C). The kinetics of fusion from the RRP and SRP were sped up by Syt-7 knockout, and this was also rescued upon reexpression (Fig. 2H). Curiously, the sustained rate was reduced in the Syt-7 KO, but this aspect was not rescued upon overexpression (Fig. 2I). However, RRP and SRP sizes in the KO + Syt-7 condition were even higher than in Syt-7 WT cells, leading to an overshoot in overall secretion within the first second after stimulation (Fig. 2F, G). Therefore, the lack of rescue of the sustained component might be caused by the twofold overexpression of Syt-7 (Fig. S2A-B), which in the presence of raised pre-stimulation [Ca^2+^] stimulates priming of vesicles beyond WT levels, moving vesicles that normally prime and fuse during the sustained component into the RRP and SRP. Recently, it was reported that kiss-and-run fusion events are upregulated in the Syt-7 KO (Zhang et al., 2019), but see (Segovia et al., 2010). Kiss-and-run fusion would cause catecholamine release without net capacitance change; we therefore performed amperometric measurements in parallel with all capacitance measurements; these measurements showed that amperometry and capacitance changed in parallel (Fig. 2A,F bottom panels; quantification of total amperometric release in Fig. S1B-C).

Almost all fast secretion from the RRP in chromaffin cells depends on Syt-1 (Nagy et al., 2006; Voets et al., 2001) (see also Fig. 1). Therefore, the data showing potentiation of the RRP size by Syt-7 indicates a cooperative/competitive interplay between Syt-7 and Syt-1, such that Syt-7 increases the amplitude of Syt-1 driven exocytosis (cooperation, Fig. 2G), but slows down its fusion kinetics (competition, Fig. 2H), at least under conditions were prestimulation [Ca^2+^] is relatively high.

The ability of Syt-7 to increase the release burst size indicates a role in vesicle priming. In order to measure Ca^2+^-dependent priming in the presence and absence of Syt-7, we varied pre-stimulation [Ca^2+^], followed by an uncaging flash to assess the secretory burst (i.e. secretion within 0.5 s after the flash, which approximately includes the RRP and the SRP). In WT cells, preflash [Ca^2+^] up to approximately 0.5 μM triggered Ca^2+^-dependent priming and a corresponding increase in burst release; at higher calcium concentrations the burst size decreases because the high prestimulation calcium causes some release before the flash (Fig. 3) (Voets, 2000). Strikingly, Ca^2+^-dependent priming was almost absent in Syt-7 KO cells (Fig. 3). In contrast, expression of Syt-7 in KO cells enhanced priming beyond WT levels at [Ca^2+^] above 0.2 μM (Fig. 3). Although Ca^2+^-dependent priming was barely detectable in the Syt-7 KO in the overall titration (Fig. 3), we noticed that the burst size was slightly higher in the Syt-7 KO in the two-group comparison (Fig. 2A vs 2F). Analysis of this showed that the RRP was significantly increased in the Syt-7 KO (low prestimulation [Ca^2+^], RRP size: 16.7 ± 3.10 fF; high prestimulation [Ca^2+^], RRP size: 29.1 ± 4.08 fF. p = 0.0480, Mann Whitney test), whereas the increase in SRP was not significant (low prestimulation [Ca^2+^], SRP size: 13.3 ± 5.35 fF; high prestimulation [Ca^2+^], SRP size: 20.7 ± 3.84 fF. p = 0.0915, Mann Whitney test). Thus, Syt-7 is necessary for strong Ca^2+^-dependent priming in adrenal chromaffin cells, and overexpressing the protein further boosts this function; in Syt-7 KO cells, a small Ca^2+^-dependent priming effect persists.

**Figure 3.**
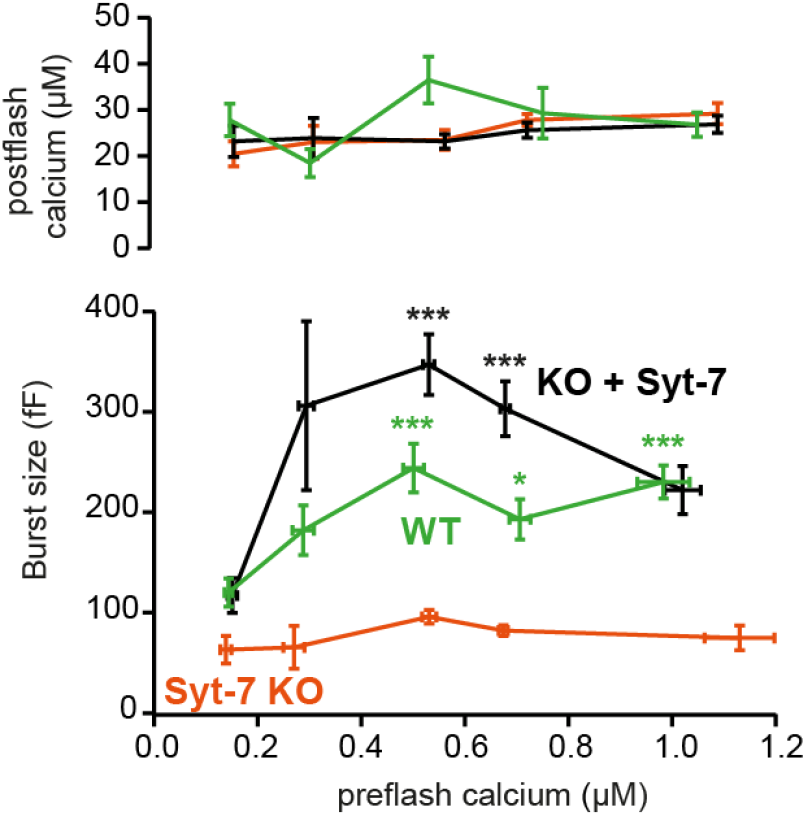
Calcium-dependent steady-state priming depends on Syt-7 expression. Titration of burst of secretion (i.e. secretion within the first 0.5 s of the uncaging flash, approximately corresponding to the fusion of the RRP and SRP) against pre-stimulation [Ca^2+^]. Top panel: post-stimulation [Ca^2+^], bottom panel: Burst size against pre-stimulation [Ca^2+^]. Syt-7 KO cells (vermilion) displayed no strong dependence on [Ca^2+^]. Calcium-dependent priming was strong in cells overexpressing Syt-7 (black), and intermediate in WT cells expressing Syt-7 at endogenous levels (green). Data information: Data are presented as mean ± SEM. *: p<0.05; *** p<0.001. Testing was by Kruskal-Wallis test with Dunn’s post-hoc test. Cells were pooled in 0.2 μM [Ca^2+^] bins from 0.0 μM to 0.8 μM and a final bin 0.8-1.2 μM for a total of 5 bins. Statistical testing is relative to the burst size at the lowest [Ca^2+^] bin for the same genotype. The number of cells in each bin from low to high [Ca^2+^]: WT: N = 29, 9, 9, 10, 22 cells; Syt-7 KO: N = 10, 12, 16, 43, 15 cells; Syt-7 KO + Syt-7: N= 12, 10, 28, 28, 20 cells.

We next investigated the role of Ca^2+^-binding to Syt-7 for its function in chromaffin cells. We mutated the ‘top loops’ of either one or both C2-domains to eliminate Ca^2+^-binding, using previously characterized substitutions of the Ca^2+^ coordinating aspartate residues (Bacaj et al., 2015). Expression analysis by immunofluorescence (Fig. S2A-B) showed that the construct mutated in the C2B domain (C2B*) was expressed at similar levels (~2-fold) as the WT Syt-7 construct. However, constructs mutated in the C2A-domain (C2A*), or both in the C2A and C2B-domain (C2AB*) were expressed at lower levels, comparable to endogenous Syt-7 levels in WT cells (Fig. S2A-B). Overexpressed protein was localized to vesicles, but some of the protein also accumulated in larger clusters (Fig. S2A). Nevertheless, data shown above (Fig. 1 and 2) demonstrated that our WT construct is functional. Using Ca^2+^ uncaging all three mutations (C2A*, C2B*, C2AB*) failed to rescue the increase in RRP and SRP size (Fig. S2C-F). Rescue by WT Syt-7 was confirmed in parallel experiments. These findings deviate from results in neurons indicating that only Ca^2+^-binding to the C2A-domain is required for Syt-7 function (Bacaj et al., 2015), which was attributed to a higher membrane-affinity of the C2A-domain (Voleti et al., 2017). However, our findings agree with a recent report that the C2 domains of Syt-7 act synergistically in binding to PI(4,5)P_2_-binding membranes (Tran et al., 2019) and previous data obtained from knockin (C2B-mutated) Syt-7 mice, which displayed a phenotype similar to Syt-7 KO in chromaffin cells (Schonn et al., 2008).

### Syt-7 reduces the depriming rate and increases the priming rate

In the above, we showed that Syt-7 is involved in LDCV priming. Priming is a reversible process (Hay and Martin, 1992; Smith et al., 1998), described by a (forward) priming rate constant *k*_*1*_, and a (backward) depriming rate constant, *k*_−*1*_ (Fig. 4A). In a simple one-pool model (without release sites, see below), the forward priming rate is [Depot] · *k*_*1*_, where [Depot] is the size of the upstream Depot pool. The steady-state RRP size in the absence of release is [Depot]·*k*_*1*_/*k*_−*1*_ (Equation 11, Materials and Methods). Under the simplifying assumption of no release during recovery, the recovery rate constant is equal to *k*_−*1*_, as recovery of the RRP is single-exponential (Equation 10, Materials and Methods):

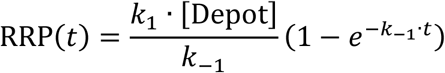

**Figure 4.**
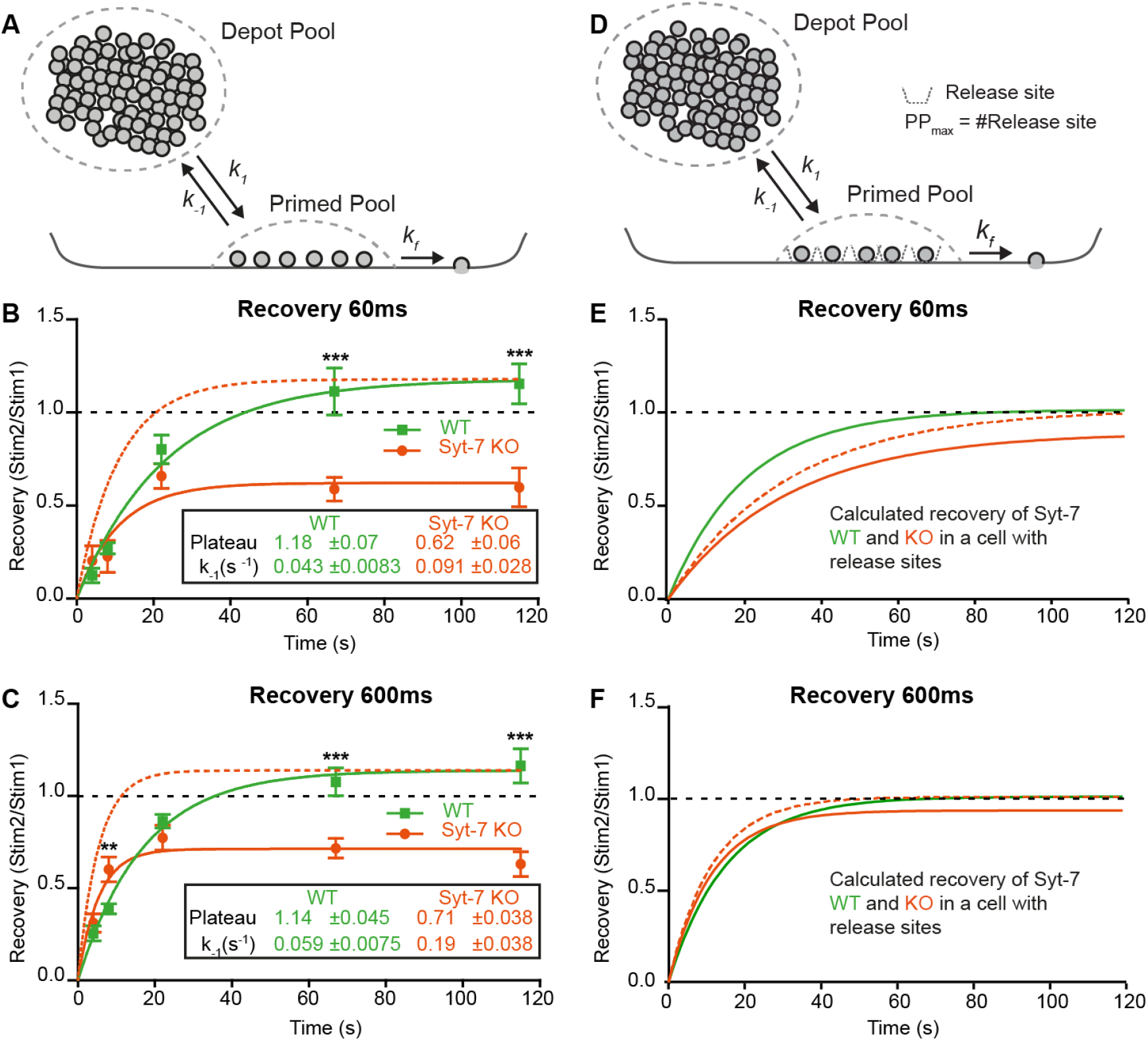
Syt-7 increases forward priming and decreases depriming. **A** Simple model (Model I) featuring a single primed vesicle pool, a reversible priming reaction (forward rate: *k*_*1*_; reverse rate: *k*_−*1*_), and a fusion rate *k*_*f*_. **B** Recovery in WT cells (green) and Syt-7 KO cells (vermilion) of secretion at 60 ms after an uncaging stimulus (see also Fig. S4), approximately corresponding to the RRP. Stim1, Stim2 = amplitude of secretion after first, or second, stimulus. Shown are mean ± SEM, plus a fit of a single-exponential recovery curve (lines). The fit to the Syt-7 KO is also shown after scaling to the same amplitude as the WT curve (vermilion broken line), to show the faster kinetics. The fitted parameters: Plateau and the rate constant of recovery, which is the rate constant for depriming, *k*_−*1*_, under simplified assumptions (see text). Both Plateau and *k*_−*1*_ are significantly different from each other (Extra sum-of-squares F test for comparison of models, p<0.0001). **C** Same as B, but secretion at 600 ms after uncaging was used, approximately corresponding to the fusion of both RRP and SRP. **D** Model (*Model II*) featuring a single primed vesicle pool, limited by a fixed number of release sites, PP_max_, a reversible priming reaction (forward rate: *k*_*1*_; reverse rate: *k*_−*1*_), and a fusion rate *k*_*f*_. **E** Recovery (at 60 ms) in the WT (green, *Model II*) with parameters recalculated from the fit of *Model I* to the data (panel B, see Methods), and Syt-7 KO curve (vermilion), with the same change in priming and depriming rate as observed experimentally, now translated to a release site model, and after scaling to the WT amplitude (vermilion broken line). Under these circumstances, recovery in the Syt-7 KO trails the WT. **F** Recovery (at 600 ms) in the WT (*Model II*) with parameters recalculated from panel C, and in the KO (vermilion), when introducing the same changes as observed experimentally, translated to a release site model, and after scaling to the WT amplitude (vermilion broken lines). Under these circumstances, the Syt-7 KO leads the WT trace, but the differences are small. Data information: Data are presented as mean ± SEM. **: p<0.01; ***: p<0.001. In (B, C) Student’s t-test: test between genotypes (WT vs. KO) at the same inter stimulus intervals; Mann Whitney test: (600ms: WT22s vs. KO22s)).

Thus, the priming rate (*k*_*1*_) only changes the steady-state pool size, but not recovery kinetics (Fig. S3A).

To distinguish between effects of Syt-7 on priming ([Depot] · *k*_*1*_) and depriming (*k*_−*1*_), we applied two sequential uncaging flashes separated by different inter-stimulus intervals in Syt-7 KO and WT cells (Fig. S4A-C). We kept the prestimulation [Ca^2+^] relatively low (250-350 nM). Due to the small amplitudes at short recovery intervals, fitting of exponentials was not reliable. Instead, we used the capacitance increase at 60 ms after the flash (approximately corresponding to fusion of the RRP), and at 600 ms (corresponding to the fusion of both RRP and SRP) to assess recovery. Recovery was clearly visible in mean capacitance traces in both the Syt-7 KO and the WT (Fig. S4A-C). Plotting the recovery curves (Fig. 4B-C), they displayed a slight overshoot in the WT case, but remained incomplete in the Syt-7 KO, even after 120 s. Fitting single exponential recovery curves allowed us to directly estimate the depriming rate, *k*_−*1*_ (see above). This rate was significantly higher in the Syt-7 KO whether the 60 ms or the 600 ms time point was considered (Fig. 4B-C, Table 1). Together with the different steady-state pool sizes, this allows us to calculate the forward priming rate, i.e. [Depot] · *k*_*1*_, which was similar for Syt-7 KO and WT at 60 ms (WT: 3.21 ± 0.63 fF/s, Syt-7 KO: 3.07 ± 0.95 fF/s; Table 1) and even higher for the Syt-7 KO than the WT at the 600 ms time point (WT: 8.60 ± 1.12 fF/s, Syt-7 KO: 15.8 ± 3.32 fF/s; Table 1). The calculated numbers are the priming rates before the first stimulation, because they derive from the pool sizes measured at the first stimulation. We interpret the slight overfilling (in the WT) or the incomplete recovery (in the Syt-7 KO) as originating from post-stimulation changes in the forward priming rate. Accordingly, in the WT the priming rate increases slightly after stimulation; however, in the Syt-7 KO the priming rate drops following stimulation (WT at 60 ms: 3.21 ± 0.63 fF/s before and 3.78 ± 0.78 fF/s after stimulation, Syt-7 KO at 60 ms: 3.07 ± 0.95 fF/s before and 1.90 ± 0.62 fF/s after secretion, Table 1). Given that the priming rate is [Depot] · *k*_*1*_, and Ca^2+^-uncaging likely leads to some reduction in [Depot], a compensatory increase in *k*_*1*_ is necessary to keep the priming rate constant from one stimulation to the next, and an even larger *k*_*1*_ increase is necessary to support overfilling in the WT. We conclude that *k*_*1*_ increases in the Syt-7 WT after stimulation, whereas in the Syt-7 KO this increase is absent or insufficient, leading to incomplete recovery.

**Table 1.** Secretion parameters for Syt-7 WT and KO. Estimated parameters for Syt-7 WT and KO when secretion is measured 60 ms or 600 ms after Ca^2+^-uncaging, which corresponds approximately to the fusion of the RRP, or the RRP + SRP, respectively. Syt-7 elimination resulted in an increase in the reverse priming rate (k_−1_), and a reduction in forward priming rate (k1*DP, where DP is the size of the Depot Pool, and k_1_ is the rate constant for priming) following stimulation.

|  | 60 ms (approx. RRP) |  | 600 ms (approx. RRP+SRP) |  |
| --- | --- | --- | --- | --- |
|  | Syt-7 WT | Syt-7 KO | Syt-7 WT | Syt-7 KO |
| **Pool size (fF)** | 74.2 ± 3.3 | 33.6 ± 0.77 | 145 ± 4 | 85.4 ± 2.7 |
| **k<sub>-1</sub> (s<sup>-1</sup>)** | 0.043 ± 0.0083 | 0.091 ± 0.028 | 0.059 ± 0.0075 | 0.19 ± 0.038 |
| **k<sub>1</sub> (Before Stim) (fF/s)** | 3.21 ± 0.63 | 3.07 ± 0.95 | 8.60 ± 1.12 | 15.8 ± 3.32 |
| **k<sub>1</sub> (After Stim) (fF/s)** | 3.78 ± 0.78 | 1.90 ± 0.62 | 9.8 ± 1.3 | 11.29 ± 2.44 |

Previously, slowed recovery kinetics in the Syt-7 KO was reported in mammalian synapses (Chen et al., 2017; Liu et al., 2014) (but see (Guan et al., 2020), for different finding in the *Drosophila* neuromuscular junction). In chromaffin cells, we found incomplete, but kinetically faster, recovery kinetics (Fig. 4B-C); we next considered how these two observations might be reconciled. In chemical synapses, priming relies on distinct release sites, which limit the size of the RRP. Specialized release sites are missing in chromaffin cells, where the RRP (and SRP) are free to change size when priming rates change – this assumption was implicit in the presentation above (*Model I*, Fig. 4A). Thus, manipulations that change priming can cause large changes in the RRP and SRP size in chromaffin cells, but not in neurons (an example is the effect of β phorbolesters (Basu et al., 2007; Lou et al., 2005; Smith et al., 1998)).

To investigate the consequences of release sites, we constructed *Model II*, where a fixed number of release sites put an upper limit on the primed pool size (Materials and Methods, Fig. S3B). The release sites were assumed to be 90% occupied at rest and to recycle immediately after use, which yields mono exponential recovery, making it possible to recalculate all parameters of *Model II* directly from our parameter estimates in *Model I* to yield recovery curves with identical kinetics. In this “neuronal” model, changing depriming (*k*_−*1*_) had much smaller effects on the recovery kinetics (Fig. S3B). Decreasing forward priming (*k*_*1*_) reduced both recovery kinetics and pool size (Fig. S3B), which is different from *Model I* (where *k*_*1*_ does not affect recovery kinetics). We finally asked how the observed changes in depriming and priming rate in the Syt-7 KO would affect recovery if translated directly to a release site model (Fig. 4D). Using recovery of the 60 ms pool (the RRP), the reduction in forward priming rate following stimulation dominated recovery, resulting in slower recovery in the Syt-7 KO than in the WT (Fig. 4E, vermilion and green curve). Considering the 600 ms pool (SRP + RRP), recovery in the Syt-7 KO preceded the WT, also after normalization (Fig. 4F), which is due to the overall higher forward priming rates we estimated for this pool in the Syt-7 KO; however, the recovery curves were overall similar, and probably indistinguishable in the presence of experimental variation.

We conclude that the faster, but incomplete recovery kinetics of the RRP in the Syt-7 KO chromaffin cell translates into slower recovery kinetics in a synapse, which reconciles our findings with data from mammalian synapses (Chen et al., 2017; Liu et al., 2014). It also demonstrates that an experimental advantage of the chromaffin cell is that effects on (forward) priming and (reverse) depriming rates can be separated - this is very difficult in the presence of a limited number of release sites.

NSF (N-ethylmaleimide sensitive factor) disassembles cis-SNARE-complexes following fusion, but it might also disassemble trans-SNARE-complexes leading to depriming, unless this activity is blocked by Munc18-1 and Munc13-1 (He et al., 2017; Ma et al., 2013; Prinslow et al., 2019). It was found that treatment with N-ethylmaleimide (NEM) stabilized the primed vesicle state in neurons where priming had been rendered labile through deletion of Munc13-1 or replacement of Munc18-1 with Munc18-2 (He et al., 2017). The effect was interpreted as the prevention of depriming through inhibition of NSF (N-ethylmaleimide sensitive factor). Hence, we asked whether Syt-7 might play a similar role in depriming. We performed Ca^2+^-uncaging in Syt-7 KO and WT cells from low pre-stimulation [Ca^2+^] (data on higher prestimulation [Ca^2+^] below) in the presence or absence of 200 μM NEM, which was included in the pipette solution. Strikingly, this led to an increase in the burst and total amount of release in the Syt-7 KO (Fig. 5A-D). Indeed, in the presence of NEM, secretion in the Syt-7 KO became indistinguishable from WT, whereas in WT cells, NEM was without significant effect (Fig. 5A-D). Kinetic analysis showed that NEM specifically increased RRP size (Fig. 5E), but with no significant effect on SRP size (Fig. 5G) or on the fast or slow time constants of release. Since no effect of NEM was found in the Syt-7 WT, the effect of NEM is occluded by Syt-7 expression. This finding is consistent with a function of Syt-7 to protect SNARE-complexes from disassembly, and it correlates well the function of Syt-7 to decrease the depriming rate. In the Syt-7 KO, SNARE-complexes are left (partly) unprotected and priming can therefore be rescued by NEM, whereas in the Syt-7 WT, where SNARE-complexes are already protected, NEM has no effect.

**Figure 5.**
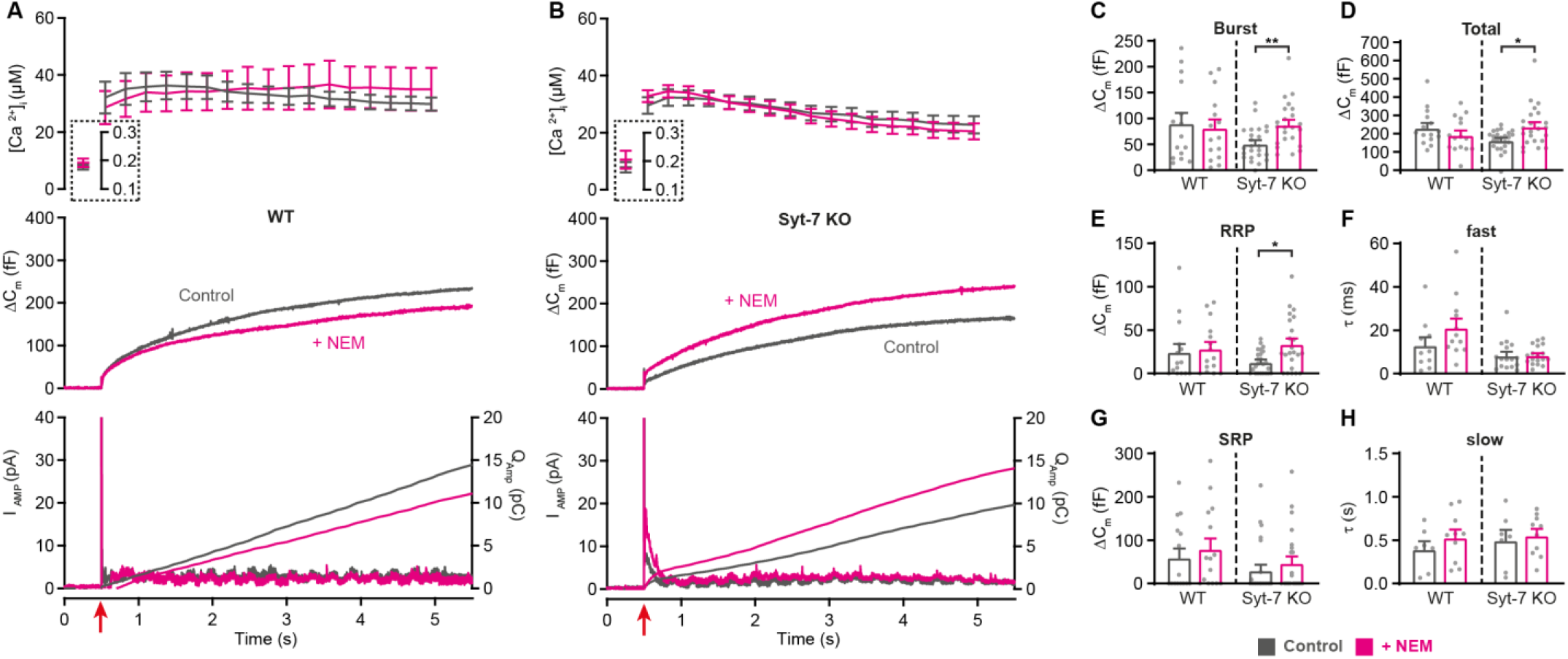
Blocking NSF-dependent de-priming occludes the effect of Syt-7 at low prestimulation Ca^2+^. **A,B** Calcium uncaging experiment from low prestimulation [Ca^2+^] in WT (A) and in Syt-7 KO (B) control cells (Control, grey) and in cells infused with 200 μM N-Ethylmaleimide (+NEM, magenta). Panels are arranged as in Fig. 1A. **C** Size of the burst (secretion within 0.5 s after the flash). **D** Total release (secretion within 5 s after the flash). **E** Size of the RRP. **F** Time constant, τ, of fusion for fast (i.e. RRP). **G** Size of the SRP. **H** Time constant, τ, of fusion for slow (i.e. SRP). Data information: Data are presented as mean ± SEM. *: p<0.05; **: p<0.01, Mann Whitney test comparing control cells to cells infused with NEM from the same genotype. Number of cells: WT control: N = 15 cells; WT + NEM: N = 15 cells; Syt-7 KO control: N = 24 cells; Syt-7 KO + NEM: N = 23 cells.

Overall, these data show that Syt-7 acts to promote priming calcium-dependently and also to inhibit depriming, likely by preventing SNARE-complex disassembly.

### Syt-7 assists in ubMunc13-2/phorbolester-dependent priming

The involvement of Syt-7 in priming prompts the question how this function relates to the canonical Munc13 priming proteins? These proteins contain the essential MUN-domain, which opens up syntaxin-1 within the syntaxin-1:Munc18-1 complex to enable SNARE-complex assembly (Basu et al., 2005; Stevens et al., 2005; Yang et al., 2015). SNARE-complex formation in turn coincides with vesicle priming (Sorensen et al., 2006; Walter et al., 2010). Thus, it is important to understand whether functional interrelationships between synaptotagmin-7s and Munc13 proteins co-determine their roles in priming. In adrenal chromaffin cells, ubMunc13-2 (= ubiquitous Munc13-2) is the dominant Munc13 isoform (Man et al., 2015), and Munc13-2 overexpression is the strongest known manipulation to increase the primed LDCV pool in chromaffin cells (Zikich et al., 2008). Secretion is also potentiated by phorbolesters (Smith et al., 1998), which activate Munc13-proteins and Protein Kinase C (Rhee et al., 2002; Wierda et al., 2007). In chromaffin cells, the effect includes a potentiation of the priming step, leading to a larger exocytotic burst (Smith et al., 1998). We investigated whether Syt-7 is involved in phorbolester- and ubMunc13-2 dependent priming.

We first tested the ability of the phorbolester phorbol-12-myristate-13-acetate (PMA, 100 nM) to increase secretion in chromaffin cells (Fig. 6) while stimulating secretion from low prestimulation [Ca^2+^]_i_ (<200 nM). Under these circumstances, PMA resulted in a robust increase in secretion in WT cells, resulting from an increase in SRP and RRP size, but not of the sustained component (overall secretion increased by 126%; Fig. 6A-D; see Fig. S1D for quantification of amperometry). PMA further led to faster SRP secretion (Fig. 6C), whereas RRP secretion kinetics were not affected. However, in Syt-7 KO cells, the situation was different: PMA resulted in only a minor potentiation of overall release (21%, Fig. 6E-H), due to a small increase in RRP (which was statistically significant) and SRP size (p=0.0506; Fig. 6F). Thus, Syt-7 is necessary for PMA to exerts its full priming effect at low prestimulation [Ca^2+^] (data on higher prestimulation [Ca^2+^] are presented below).

**Figure 6.**
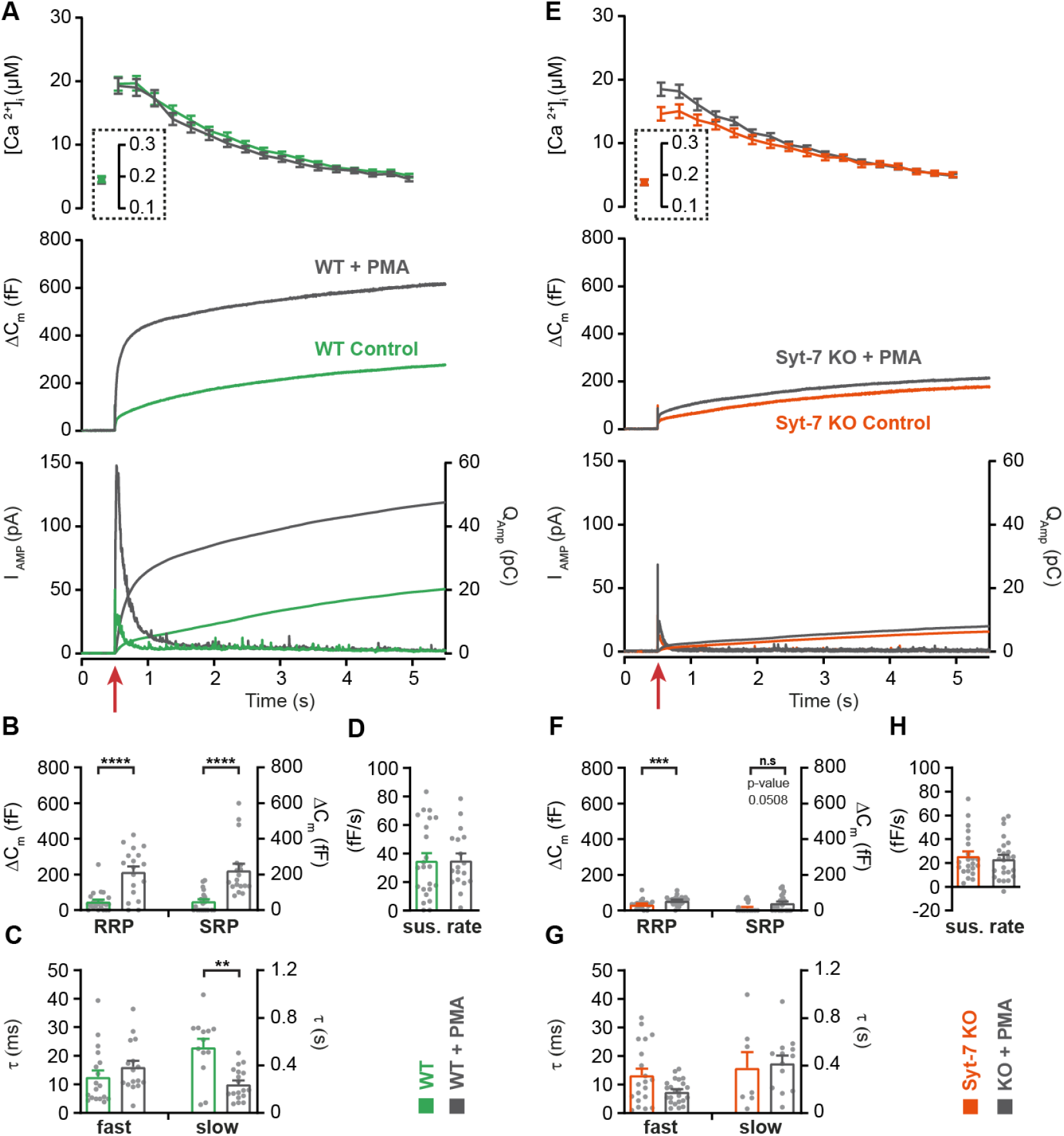
Syt-7 stimulates PMA-induced potentiation of release at low prestimulation Ca^2+^. **A** Calcium uncaging experiment from low prestimulation [Ca^2+^] in Syt-7 WT cells (green) and in Syt-7 WT cells perfused with 100 nM phorbol 12-myristate 13-acetate (PMA) (WT + PMA) (grey). Panels are arranged as in Fig. 1A. PMA treatment strongly augmented the primed pool size in WT cells. **B** Sizes of the RRP and SRP. **C** Time constants, τ, of fusion for fast (i.e. RRP) and slow (i.e. SRP) secretion. **D** Sustained rates of secretion. **E** Calcium uncaging experiment from low prestimulation [Ca^2+^] in Syt-7 KO cells (vermilion) and in Syt-7 KO cells perfused with 100 nM PMA (KO + PMA) (grey). PMA-induced potentiation of release was much weaker in Syt-7 KO cells. **F** Size of the RRP and SRP. **G** Time constants, τ, of fusion for fast (i.e. RRP) and slow (i.e. SRP) secretion. **H** Sustained rate of secretion. Data information: Data with error bars (A-H) are presented as mean ± SEM; in (A, E), the traces are the average of all cells. Statistics: *: p<0.05; ** p<0.01; *** p<0.001; **** p<0.0001. Analysis was performed with Mann-Whitney test. Number of cells: WT: N = 23 cells; WT + PMA: N = 18 cells; Syt-7 KO: N = 22 cells; Syt-7 KO + PMA = 24 cells.

Overexpression of ubMunc13-2 in WT chromaffin cells using a Semliki Forest Virus construct resulted in massive secretion, which reached 1.0-1.5 pF (Fig. 7A; data from Syt-7 WT and KO are the same as in Fig. 2) when again assayed from a low prestimulation [Ca^2+^] (data on higher prestimulation [Ca^2+^] below). This was due to a massive increase in both RRP and SRP sizes (Fig. 7D, G), as previously reported from bovine chromaffin cells (Zikich et al., 2008). However, we noted that the capacitance curve of the ubMunc13-2 overexpression cells did not have the normal concave form; instead, the curves showed signs of a secondary acceleration, giving rise to a convex curve - this was especially clear in the case of ubMunc13-2 expressed in Syt-7 KO cells (Fig. 7A-C). This change in secretory kinetics was also observed in the parallel amperometric measurements (Fig. 7A bottom), where the amperometry current increased again after the first phase, reaching a second maximum around 1.5 s. An inspection of individual traces revealed that the SRP was fusing after a longer delay, whereas RRP fusion kinetics were largely unaffected (Fig. 7B-C shows examples). To investigate whether this is a feasible interpretation, we fitted secretory traces with an alternative function for a SRP fusing after a delay. The function describing SRP fusion is

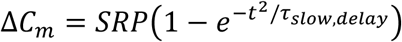

where *τ*_*slow,delay*_, is the product of the time constants for the delay and for slow fusion itself. Due to the long delay, it is not possible to determine fusion kinetics and delay separately (see Materials and Methods for derivation of this function). We used the chi-square value between fit and data to determine whether this function or the standard sum of two exponentials and a line (see Materials and Methods) fitted the traces better. Syt-7 KO cells overexpressing ubMunc13-2 could only be fitted with the delayed-SRP model (yielding the best fit in 16 of 17 cells), but also Syt-7 WT expressing ubMunc13-2 more often than not had a delayed SRP (best fit in 10 out of 17 cells). This difference was statistical significant (p=0.0391, when tested as a contingency table using Fisher’s exact test). This indicates that after overexpression of ubMunc13-2 the SRP fuses after a delay in the absence of Syt-7, whereas in the presence of Syt-7 a delay is sometimes present, but not always. In Fig. 7A-G, we compare the results after ubMunc13-2 expression in WT and Syt-7 KO to the data obtained from WT and Syt-7 KO without overexpression (stimulated from a low prestimulation [Ca^2+^], Fig. 2A-D). Kinetic analyses are presented in Fig. 7D-H (note that cells from WT and Syt-7 KO were fitted with 2 exponentials and a line, Eq. 1, and not obtained in parallel with the other data; therefore these data were not statistically compared to ubMunc13-2 overexpressing cells). These data show a larger ability of ubMunc13-2 to increase the RRP in Syt-7 WT compared to Syt-7 KO (Fig. 7D), whereas ubMunc13-2 overexpression increased the SRP-size both in the presence and absence of Syt-7 (Fig. 7G).

**Figure 7.**
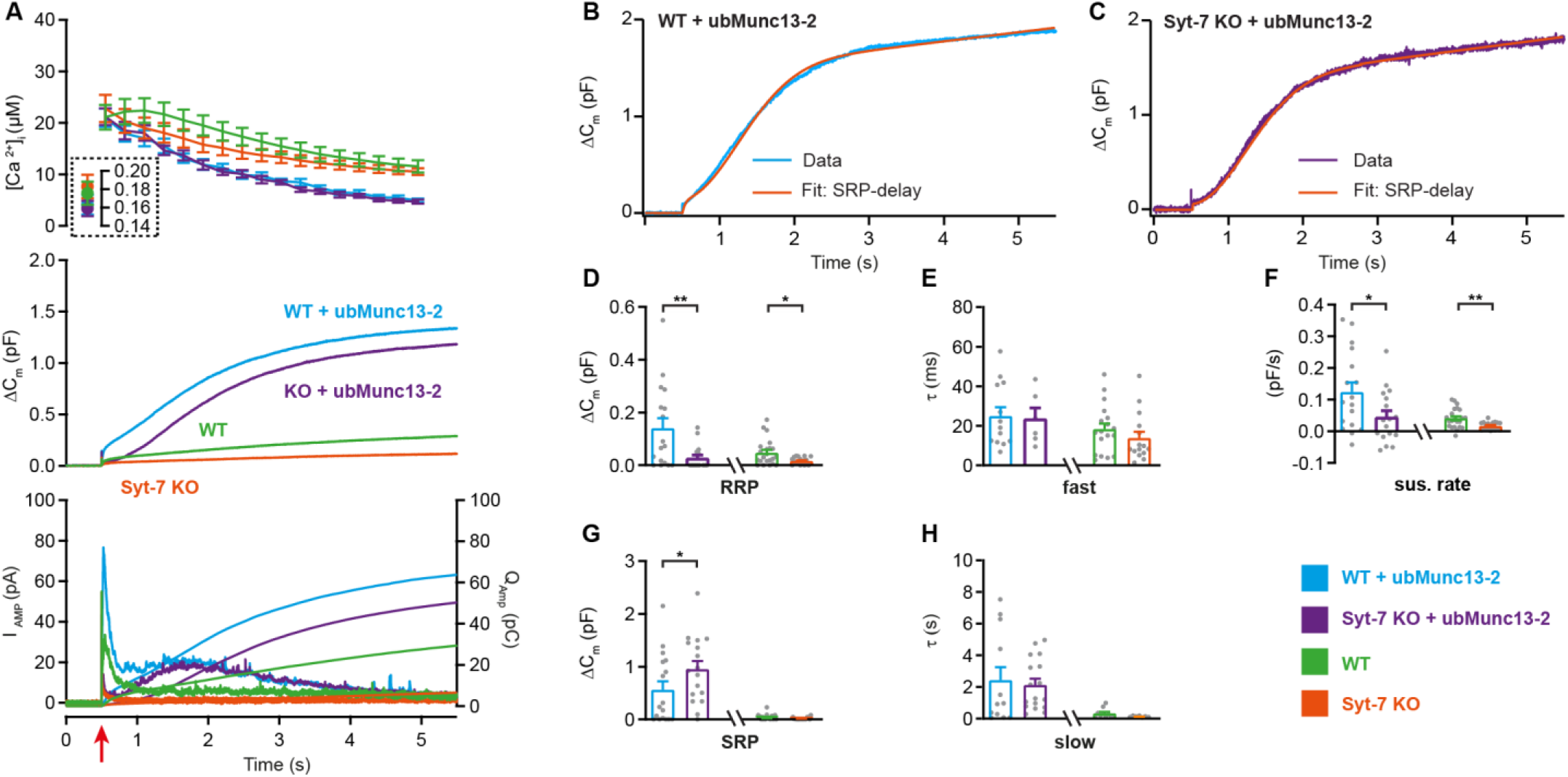
Syt-7 stimulates ubMunc13-2 dependent priming at low prestimulation Ca^2+^. **A** Calcium uncaging experiment in WT overexpressing ubMunc13-2 (WT + ubMunc13-2) (cyan traces), and Syt-7 KO overexpressing ubMunc13-2 (KO + ubMunc13-2) (purple traces) stimulated from a low prestimulation [Ca^2+^]. Data from Syt-7 WT and Syt-7 KO are the same as in Fig. 2A. Panels are arranged as in Fig. 1A. The overexpression of ubMunc13-2 potentiated the release in Syt-7 KO cells after a remarkable delay of the SRP. **B** An example capacitance trace (‘Data’, blue trace) from a WT cell overexpressing ubMunc13-2 (cyan trace) with a function taking into account the SRP-delay (‘Fit’, red trace). **C** An example capacitance trace (‘Data’) from a Syt-7 KO cell overexpressing ubMunc13-2 (‘Fit’, purple trace) fitted with a function taking into account the SRP-delay (red trace). **D-H** In the Syt-7 WT + ubMunc13-2 or Syt-7 KO + ubMunc13-2, for each cell recorded, the chi-square values between fit and data were used to judge whether the standard sum of two exponentials and a line function or the function including the SRP delay (see Materials and Methods) fitted the traces better. Values from the best fit were averaged to obtain the RRP, SRP sizes and time constants. **D, G** Sizes of the RRP and SRP. **E, H** Time constant, τ, of fusion for fast (i.e. RRP) and slow (i.e. SRP) secretion. Note that the τ for the SRP in the Syt-7 WT + ubMunc13-2 and Syt-7 KO + ubMunc13-2 groups include both the secretory delay and the fusion kinetics (Eq. 7). Due to the slower τ for the SRP in this data set, we assumed that a τ originated from the RRP if τ ≤ 60 ms and from the SRP when 60 ms ≤τ ≤ 1600 ms (se Materials and Methods). **F** Sustained rate of secretion. Note that in some cases when fitting with the function taking into account the SRP-delay, a negative sustained rate resulted from the fit. Data information: In (A-F) data with error bars are presented as mean ± SEM; in (A), the traces are the mean of all cells. WT (green) and Syt-7 KO (vermilion) were not obtained in parallel experiments, but are displayed here only to illustrate the increase upon ubMunc13-2 overexpression; statistical tests are only conducted for Syt-7 KO + ubMunc13-2 vs Syt-7 WT + ubMunc13-2 and Syt-7 WT vs Syt-7 KO. Note that the RRP size of WT vs Syt-7 KO is significantly differient, due to the two-group comparison, which was not the case in Fig. 2A (three-group comparison). Statistics: *: p<0.05; **: p<0.01, Mann Whitney test. Number of cells: WT + ubMunc13-2: N = 17; Syt-7 KO + ubMunc13-2: N = 17 cells; WT: N = 22; Syt-7 KO: N = 19.

Overall, we have shown that under these circumstances (low prestimulation [Ca^2+^]), PMA requires Syt-7 to strongly increase RRP and SRP size, whereas Munc13-2 requires Syt-7 to increase the RRP, but not the SRP size. However, the SRP tends to fuse with an additional delay in the absence of Syt-7.

### Synaptotagmin-7 independent mechanisms partly stimulate priming at high prestimulation [Ca^2+^]

We next repeated the experiments with phorbolester, ubMunc13-2 overexpression, and NEM treatment from higher prestimulation [Ca^2+^]. These data are presented in Fig. S5, S6, S7. At higher prestimulation [Ca^2+^]_i_, phorbolester led to a strong increase in RRP size in Syt-7 WT cells, no significant changes in SRP and a mild, but not significant, increase in the sustained component (Fig. S5A-D). In Syt-7 KO, the RRP was also potentiated by PMA (Fig. S5F), although the RRP size in the Syt-7 KO remained smaller than in the Syt-7 WT (Fig. S5B). In addition, the sustained component was significantly potentiated by PMA in the Syt-7 KO. Overall, therefore, total secretion in the Syt-7 KO + phorbolester was only slightly reduced compared to Syt-7 WT + phorbolester (see also Fig. S1E), but the size of the RRP was smaller and secretion in the absence of Syt-7 was therefore overall slower.

When expressing ubMunc13-2 in Syt-7 KO and Syt-7 WT and stimulating from a higher prestimulation [Ca^2+^], the RRP was clearly increased by ubMunc13-2 in the Syt-7 KO, although still depressed compared to ubMunc13-2 overexpressing Syt-7 WT cells (ubMunc13-2 in Syt-7 KO: 410 ± 83 fF; ubMunc13-2 in Syt-7 WT: 207 ± 39 fF, p=0.04; Fig. S6D). Moreover, in ubMunc13-2 expressing WT cells, the two exponentials and a line fitted 12 of 13 cells best, and only one cell was fitted better with a SRP-delay. In contrast, in Syt-7 KO cells the function incorporating a SRP-delay fitted 16 of 16 cells better. This difference was statistical significant (p<0.0001, when tested as a contingency table using Fisher’s exact test). This demonstrates that fusion of the ubMunc13-2-induced SRP still occurs in the absence of Syt-7, but that it does so with a significant delay compared to Syt-7 WT cells.

Finally, neither Syt-7 KO nor WT cells were affected by NEM at high pre-stimulation [Ca^2+^](Fig. S7), thereby indicating that alternative molecules protect SNARE complexes under these conditions. These data are consistent with two alternative Syt-7 functions, as demonstrated above, namely the promotion of forward priming (*k*_*1*_) and the inhibition of unpriming (*k*_−*1*_). Syt-7 might assist in SNARE-complex formation to increase *k*_*1*_ at high prestimulation [Ca^2+^], and then protect the formed SNARE-complexes (reducing *k*_−*1*_) as the [Ca^2+^] relaxes back to low values, however, Syt-7 does not subserve both functions simultaneously and alternative factors appear responsible for protecting SNARE-complexes at high [Ca^2+^].

Altogether, these data show that when combining prestimulation [Ca^2+^] with additional manipulations to increase priming, some priming is seen in the absence of Syt-7, although the RRP size often remains smaller. These data can be accounted for by a mechanism in which Syt-7 priming acts upstream of Munc13-2/phorbolester (see Discussion).

### Syt-7 promotes the placement of dense-core vesicles at the plasma membrane

High-pressure freezing (HPF) fixation followed by freeze-substitution for electron microscopy has become an established approach for the analysis of secretory vesicle docking in systems such as hippocampal organotypic slice cultures (Imig et al., 2014; Siksou et al., 2009), and acute adrenal slices (Man et al., 2015). This method, especially when combined with high-resolution 3D electron tomography (3D-ET), allows an accurate assessment of SV and LDCV placement down to a few nanometers of distance to the plasma membrane. The assembled SNARE-bundle can bridge membranes separated by as much as 9-15 nm in the presence of complexin (Li et al., 2011). Therefore, high-resolution 3D-ET makes it possible to distinguish the close apposition of vesicle and membrane that follows from the assembly of the priming complex, yielding a morphological read-out of priming (Imig et al., 2014; Siksou et al., 2009). Previous work showed no difference in placement of LDCVs upon deletion of Munc13-1 and Munc13-2 (Man et al., 2015). As Syt-7 partly acts to limit depriming at low resting [Ca^2+^] (above), HPF might offer a possibility of correlating LDCV priming with morphological changes.

Following previously established protocols (Man et al., 2015), we combined HPF and freeze-substitution with classical 2D-EM and high-resolution 3D-ET. The reason for including 2D-EM is to quantify the overall vesicle distribution and vesicle number in the entire cell, whereas 3D-ET focuses only on membrane-proximal vesicles. Quantitative analysis of 2D-EM images of Syt-7 WT and KO chromaffin cells (Fig. 8A-D) revealed no differences in vesicle distribution within 2 μm of the plasma membrane (Fig. 8E-F), in the number of LDCVs per cell profile (Fig. 8G), the density of LDCVs per cytoplasm area (Fig. 8H), or in the fraction of LDCVs present within 40 nm of the membrane (“membrane-proximal” vesicles; Fig. 8I). Close LDCV-membrane apposition was analyzed using 3D-ET. Focusing exclusively on LDCVs placed less than 100 nm from the membrane, the proportion of vesicles placed closer than 40 nm was not changed in the Syt-7 KO (Fig. 8S). Vesicles observed to be in physical contact with the plasma membrane in tomographic volumes were considered ‘docked’ and based on the voxel dimension of reconstructed tomograms these vesicles were placed in the 0-4 nm bin (Man et al., 2015). The number of docked vesicles (0-4 nm) and vesicles placed between 4 and 6 nm from the plasma membrane was reduced in the Syt-7 KO (Fig. 8P); only the latter difference was statistically significant. When pooled into a single bin, the number of vesicles placed closer than 6 nm from the plasma membrane was significantly reduced in the Syt-7 KO (Fig. 8T, p = 0.018). The lack of significance in the docked bin (0-4 nm) might be due to a large number of ‘dead-end’ docked vesicles (Hugo et al., 2013; Verhage and Sorensen, 2008). Interestingly, the reduction of LDCVs in the vicinity of the membrane in Syt-7 KO cells was accompanied by an significantly increased number of vesicles at slightly larger distances (20-40 nm, insert in Fig. 8P). We also noted a tendency for vesicles to be of slightly smaller diameter in the Syt-7 KO (Fig. 8R). Quantitative considerations (Materials and Methods) combining the overall density of vesicles (as observed in 2D images) with the accurate determination of vesicle diameter (as obtained in 3D-ET) made it possible to estimate the total number of vesicles in Syt-7 WT cells (~12,334 vesicles) and in Syt-7 KO cells (~13,270 vesicles), whereas the total number of vesicles that are attached to the plasma membrane (bin 0-4 nm) was ~329 vesicles/cell for Syt-7 WT and ~266 vesicles/cell for Syt-7 KO.

**Figure 8.**
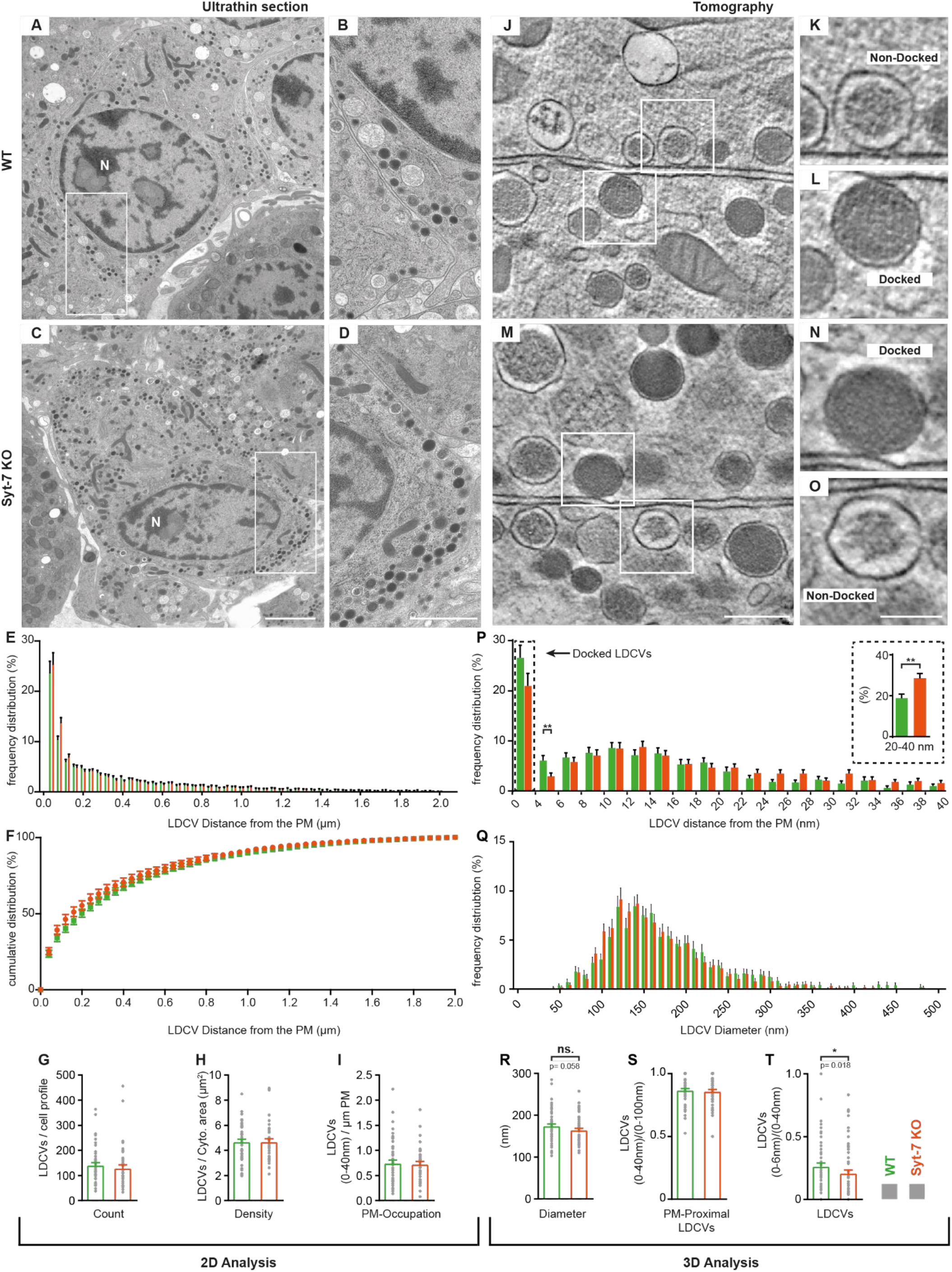
Syt-7 induces membrane-apposition of LDCVs. **A, C** 2D-EM micrographs of ultrathin adrenal slices from WT (A) and Syt-7 KO (C) newborn mice. Nucleus is designated (N). Scale bar: 2 μm. **B, D** Magnification of selection in (A) and (C), respectively. Scale bar: 1 μm. **E** Frequency distribution of large dense core vesicles (LDCVs) within 2 μm from the plasma membrane (PM) in WT (green) and Syt-7 KO (vermilion) cells. **F** Cumulative frequency plot of (E). **G** Total number of LDCV per cell profile. **H** Number of LDCVs per cytosolic area (Density = LDCVs/μm^2^). **I** Number of PM-Proximal LDCVs per μm of PM circumference. 2D analysis revealed normal cell morphology and LDCV distribution in the absence of Syt-7. **J, M** 3D-EM reconstructed tomogram subvolume from WT (J) and Syt-7 KO (M) showing two cells with opposing membranes. Scale bar: 300 nm. **K, L, N, O** Magnifications of selected regions in (J) and (M); showing docked (L, N) and non-docked (K, O) LDCVs. Scale bar: 150 nm **P** Frequency distribution of PM-proximal LDCVs where docked vesicles are accumulated in the 0-4 nm bin. Insert is a summation of 20-40 nm into single bins. **Q** Frequency distribution of LDCV diameter. **R** Diameter of LDCVs within 100 nm from the PM. **S** PM-Proximal LDCVs (0-40 nm) normalized to (0-100 nm) LDCVs (0-40 nm LDCVs/0-100 nm LDCVs). **T** Vesicles within 6 nm of the membrane (bin 0-6 nm) normalized to PM-Proximal LDCVs (0-6 nm LDCVs/0-40 nm LDCVs). Overall, the 3D analysis showed that LDCVs 0-6 nm from the PM are markedly reduced in the absence of Syt-7. Non-docked LDCVs in the Syt-7 KO accumulated at 20-40 nm from the PM. Data information: Values are mean ± SEM.*: p<0.05; **: p<0.01. Student’s t-test: (G): p = 0.4240; (H): p = 0.9711; (I): p = 0.9241; (Insert in (P): 20-40 nm): p = 0.0025; (R): p = 0.0581; (S): p = 0.6127. Mann Whitney test: (P): 4-6 nm: p = 0.0039; (T): p = 0.0184. Number of cells, 2D analysis: (WT) N = 60 cells, (Syt-7 KO) N = 46 cells; 3D analysis: (WT) N = 74 cells, (Syt-7 KO) N = 74 cells.

Overall, in the absence of Syt-7 less vesicles are placed very near the plasma membrane, whereas more vesicles become placed at distances (20-40 nm) which are probably beyond the formation range of the priming-complex. This phenotype correlates with the ability of Syt-7 to stimulate priming.

### Limited colocalization of Syt-1 and Syt-7

The work performed above identified several conditions under which Syt-7 stimulated the size of the RRP. This was the case upon elevation of intracellular calcium using flash photolysis (Fig. 2), upon stimulation by phorbolester (Fig. 5), after ubMunc13-2 expression (Fig. 6), or when treatment by NEM was performed at low prestimulation [Ca^2+^] (Fig. 7). The RRP fuses fast (time constant typically 10-20 ms at 20 μM Ca^2+^) and depends on Syt-1 expression, as shown in Syt-1 knockout and rescue experiments (Nagy et al., 2006; Voets et al., 2001), and as demonstrated here by Syt-1 expression in the Syt-1/Syt-7 Double KO cells (Fig. 1). Upon Syt-7 expression in the Syt1/Syt-7 DKO there was a minor increase in the RRP (Fig. 1); however, the time constant was increased to 50 ms, which does not correspond to a typical RRP time constant. These data indicate that Syt-7 is - in itself - not able to fuse vesicles with fast kinetics; instead, Syt-1 must fuse these vesicles. Thus, Syt-7 and Syt-1 cooperate (Walter et al., 2011), such that Syt-7 builds up a larger RRP, which then fuses with the help of Syt-1.

This conclusion raises the question whether Syt-1 and Syt-7 are localized to the same vesicles as a prerequisite for their cooperation, or whether cooperation takes place in a different way. Previous work emphasized the difference in localization between Syt-1 and −7 in PC12-cells, and in rat and bovine chromaffin cells, since only limited overlap between either endogenously or exogenously expressed Syt-1 and Syt-7 was identified in immunolabelling experiments performed at light and electron microscopic levels (Matsuoka et al., 2011; Rao et al., 2014; Wang et al., 2005; Zhang et al., 2011). In a recent study performed in mouse chromaffin cells, only 3-5% overlap between endogenous Syt-1 and Syt-7 was reported (Bendahmane et al., 2020).

We performed double immunostaining of cultured adrenal chromaffin cells from newborn mice. Both Syt-1 and Syt-7 overlapped with a marker for secretory granules (chromogranin A; Fig. S8; Manders’ coefficients: M1, Syt-1 fraction in CgA: 0.50 ± 0.05, Syt-7 fraction in CgA: 0.42 ± 0.05; M2, CgA fraction in Syt-1: 0.41 ± 0.05, CgA fraction in Syt-7: 0.47 ± 0.04). These data are consistent with previous findings that Syt-7 localizes partly to secretory granules, and partly to the endo-lysosomal system (Arantes and Andrews, 2006; Bendahmane et al., 2020).

We next performed double immunostaining using a mouse monoclonal Syt-7 antibody and a rabbit polyclonal Syt-1 rabbit antibody (both from Synaptic Systems, see Materials and Methods; Fig. 9A). In chromaffin cells from the Syt-7 KO mouse (Maximov et al., 2008) staining against Syt-7 was strongly depressed, as expected (Fig. 9B). Antisera specificity was verified in Western blots from Syt-7 KO mice (Fig. S9G), which indicated that any residual immunolabelling in Syt-7 KO cells is unspecific. Staining for Syt-1 was unchanged in the Syt-7 KO (Fig. 9B; the Syt-1 antibody was previously verified on Syt-1 KO chromaffin cells (Nagy et al., 2006)). Some co-localization between Syt-1 and Syt-7 was observed in these stainings (Fig. 9C), with Manders’ coefficients of 0.19 ± 0.02 (Syt-1 fraction in Syt-7) and 0.16 ± 0.02 (Syt-7 fraction in Syt-1). These numbers were reduced to 0.006 ± 0.003 and 0.039 ± 0.009 in the Syt-7 KO, indicating specific overlap between Syt-1 and Syt-7 signals. In these stainings, we included a small amount (0.2%) of glutaraldehyde, because we found that this branched aldehyde led to a better recognizable vesicular staining, which was a prerequisite for doing single-vesicle imaging (see below). Sodium Borohydride was used as a quenching agent. To test that glutaraldehyde does not add unspecific fluorescence, we performed stainings without glutaraldehyde, which yielded the same level of background staining (Fig. S9A-C). Another antibody combination - a mouse monoclonal Syt-7 antibody (clone 275/14, Sigma-Aldrich) together with a rabbit polyclonal Syt-1 antibody (W855, a gift from T.C. Südhof, previously verified on Syt-1 KO chromaffin cells (Kedar et al., 2015)) yielded similar results (Fig. S9D-F), but slightly lower background staining and slightly higher Manders’ coefficients (0.37 ± 0.03 for Syt-1 fraction in Syt-7; 0.24 ± 0.03 for Syt-7 fraction in Syt-1).

**Figure 9.**
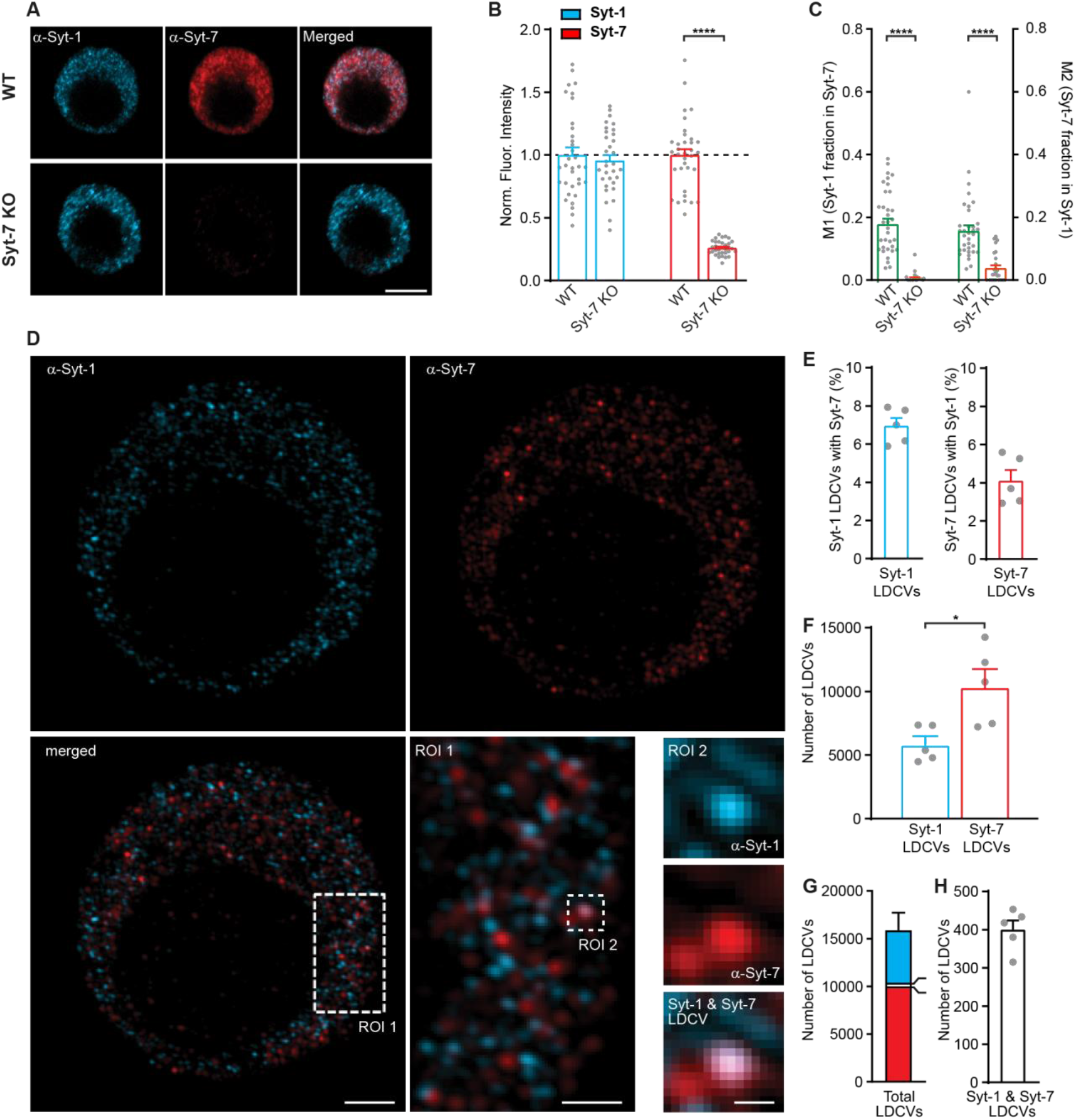
Syt-1 and Syt-7 displays limited colocalization. **A** Single confocal slices of new-born mouse chromaffin cells stained against Syt-1 (α-Syt-1) and Syt-7 (α-Syt-7) in WT cells and in Syt-7 KO cells, and merged images. Scale bar: 5 μm. **B** Quantification of staining against Syt-1 and Syt-7 in WT and Syt-7 KO cells. For staining with other antibodies: see Fig. S9D-F. **C** Manders’ coefficients M1 and M2 (mean ± SEM) for co-localization analysis of Syt-1 and Syt-7 in WT and Syt-7 KO cells. **D** Single optical slices of new-born WT mouse chromaffin cells stained against Syt-1 (α-Syt-1) and Syt-7 (α-Syt-7) acquired with 3D-structured illumination microscopy (3D-SIM). Scale bar: 2 μm. Bottom right: Magnified ROIs. ROI 1: A section of the cell from the merged-channel image showing that the majority of the vesicles are identified either as Syt-1 -or Syt-7-vesicles and few are positive for both isoforms. Scale bar: 1 μm. ROI 2: An example of a vesicle where Syt-1 and Syt-7 appear co-localized to the same vesicle (top panel: α-Syt-1; middle panel: α-Syt-7; bottom panel: A merged image of α-Syt-1 and α-Syt-7 channels). Scale bar: 0.2 μm. **E** Quantification of the percentage of Syt-1 positive vesicles that were co-stained for Syt-7 and vice versa. **F** Number of Syt-1 and Syt-7 vesicles per cell. **G** The total number of vesicles per cell. Bar colors indicate the proportions of Syt-1 (cyan), Syt-7 (red) and Syt-1/Syt-7 (white) vesicles. **H** Number of Syt-1/Syt-7 vesicles where the two isoforms are considered to be co-localized to the same vesicle as shown in ROI 2 example (D). Data information: Values are mean ± SEM. *: p<0.05; ****: p<0.0001, Student’s *t*-test. Number of cells in (B, C): Syt-7 WT: N = 22 cells; Syt-7 KO: N = 22 cells. Number of cells in (E-H): Syt-7 WT, N = 5 cells.

3D-SIM has a resolution approx. 2-fold higher than confocal microscopy, which made it possible to distinguish single objects/vesicles in both the Syt-1 and the Syt-7 channels in WT cells (Fig. 9D shows 110 nm thick optical sections). Strikingly, most Syt-1 and Syt-7 positive vesicle-like structures did not display detectable colocalization (Fig. 9D: ROI 1), but colocalization was also detected (Fig. 9D, ROI 2). The average size (full width at half maximum) of Syt-1 vesicles was 130 nm and of Syt-7 was 149 nm (note that Syt-7 was imaged at longer wavelengths, which might account for the difference). We used an automated routine to analyze every third optical section, such that we avoided detecting the same vesicles twice. We found that among the Syt-1 positive structures, 7.0 ± 0.4 % colocalized with Syt-7, whereas from the Syt-7 positive structures, 4.1 ± 0.6 % colocalized with Syt-1 (Fig. 9E). The total number of vesicle-like structures positive for Syt-1 was 5,870 ± 620 per cell; and the number of structures positive for Syt-7 10,390 ± 1,365 per cell (Fig. 9F). The number of vesicle-like structures positive for both Syt-1 and Syt-7 was 400 ± 24 (fig. 9G-H).

Even with the higher resolution of 3D-SIM compared to confocal microscopy, it cannot be ruled out that the occasionally identified co-localized structures are really two separate structures, which by chance are too close to each other to be individually resolved. To rule this out, we performed another analysis, by averaging around Syt-1 positive vesicular structures in confocal images (Walter et al., 2014). This was performed using confocal images, because the reconstruction algorithm of the 3D-SIM might complicate interpretations. By automatically detecting intensity peaks in the Syt-1 channel we could use averaging in a small Region of Interest (ROI) in both the Syt-1 and the aligned Syt-7 channel to detect a co-localized Syt-7 signal. This resulted in a clear Syt-1 positive peak (Fig. S10A), as expected, but also the averaged Syt-7 staining displayed a noticeable peak around the center (Fig. S10A). When examining a line profile through the peak, the Syt-7 signal was of much smaller magnitude (less than 10% of the total staining amplitude) than the Syt-1 peak (Fig. S10B). Importantly, when performing the same analysis in Syt-7 KO cells, the peak in the Syt-7 channel was absent (Fig. S10B). These data confirm the existence of a specific, but limited, colocalization between Syt-1 and Syt-7 positive signals (see Discussion).

## Discussion

### Functional interactions between poorly colocalized synaptotagmins

By expressing either Syt-1 or Syt-7 in Syt-1/Syt-7 DKO cells we showed that both synaptotagmins are able to act as stand-alone sensors, with Syt-7 being a slower sensor than Syt-1 both on the population level, and at the level of single LDCV fusion (Fig. 1). Isoform-specific stainings performed by others (Bendahmane et al., 2020) and us, indicate that the overlap between endogenous Syt-1 and Syt-7 positive structures is limited. Together, these data draw a deceptively simple picture: that the RRP and the SRP are equipped with separate Ca^2+^-sensor, and fuse independently.

However, further work made it clear that this picture is insufficient, as the two sensors interact. In flash photolysis experiments Syt-7 KO displayed faster fusion kinetics for both RRP and SRP fusion (Fig. 2H) than the Syt-7 WT, and this effect was rescued by Syt-7 overexpression, indicating that the slow kinetics of Syt-7 driven fusion partly affects the fusion kinetics of both pools. In addition, Syt-7 stimulates Ca^2+^-dependent priming of the RRP (Fig. 3) and assists ubMunc13-2 (Fig. 7) and PMA (Fig. 6) in stimulating the RRP size (see also below). RRP fusion requires syt-1 (Nagy et al., 2006; Voets et al., 2001). Thus, the two sensors act competitively (in setting fusion speeds) and cooperatively (in vesicle priming), which is hard to reconcile with the simple case of two independent vesicle pools. Indeed, refilling of the RRP happens at the expense of the SRP (Voets et al., 1999), and mutation in SNAREs are best explained by both slow and fast fusion along a single pathway (Walter et al., 2013). Importantly, our findings of competition and cooperation could be made relying entirely on endogenously expressed Syt-1 and Syt-7, by comparing Syt-7 WT to Syt-7 KO cells. This is important, because overexpression of Syt-7 could cause ‘overflow’ and targeting to other vesicle populations; nevertheless, overexpression overall rescued Syt-7 KO secretion and Ca^2+^-dependent priming (Fig. 2-3).

The evidence indicating functional interaction between Syt-1 and Syt-7 contrasts with the limited co-localization of the two sensors. Limited colocalization in spite of functional interaction is also found in neurons, where Syt-7 was described as a plasma membrane sensor (Sugita et al., 2001), and imaging of pHluorin-tagged Syt-7 showed almost no or delayed exocytosis (Dean et al., 2012; Li et al., 2017; Weber et al., 2014), except at higher stimulation strengths, where plasma membrane localized syt-7-pHluorin was initially endocytosed, and then recycled back to the plasma membrane (Li et al., 2017). Syt-7 is a prerequisite for synaptic facilitation (Jackman et al., 2016; Turecek et al., 2017), which involves stimulation of fast release, which implies cooperation between Syt-7 and the fast sensors (Syt-1/Syt-2/Syt-9 (Xu et al., 2007) - note different findings in *Drosophila* neuromuscular junction (Guan et al., 2020)).

Either Syt-1 and Syt-7 cooperate using an indirect mechanism, or else limited colocalization – beyond the resolution of most methods - allows for their functional interaction. Indirect cooperation could come about by fusion of Syt-7 positive vesicles at calcium concentrations <1 μM, which would allow Syt-7 to interact with Syt-1 positive vesicles from the plasma membrane side subsequently, causing their priming (“cross-priming” of vesicles). Indeed, TIRF microscopy has demonstrated that the higher affinity of Syt-7 confers vesicles with the ability to fuse at lower calcium concentrations than Syt-1 (Rao et al., 2014; Rao et al., 2017; Wang et al., 2005). This model is in agreement with the ability of both Syt-1 and Syt-7 to act as stand-alone Ca^2+^-sensors for exocytosis. However, some colocalization between Syt-1 and Syt-7 may exist in chromaffin cells. We showed using an averaging routine that single Syt-1 positive structures contain a small, but specific, Syt-7 positive signal (Fig. S10). Thus Syt-1 and Syt-7 might co-localize on some or even most vesicles. In this case, vesicles might prime through interaction of Syt-7 with the plasma membrane, SNAREs, and other priming factors, whereas fusion could be executed partly by Syt-1 and partly by Syt-7, with the overall fusion kinetics being determined by both. This model accounts for both the competitive and synergistic interaction between the synaptotagmins.

### Syt-7 in Ca^2+^-dependent priming of fast release

The presence of Syt-7 at endogenous levels caused a larger RRP and SRP size [here and (Schonn et al., 2008)] - and 2-fold Syt-7 overexpression increased RRP (and SRP) size even more (Fig. 2G). Titration of the burst (RRP+SRP) as a function of prestimulation [Ca^2+^] revealed strong Ca^2+^-dependent priming in the presence of Syt-7, but not in its absence (Fig. 3).

A function for Syt-7 in speeding up RRP replenishment in neurons was reported by (Liu et al., 2014), whereas Bacaj et al. (2015) found that Syt-7 helps maintain the size of the RRP. Intuitively one would expect the first experiment to be due to a difference in priming rate, whereas the latter would seem to imply an effect de-priming rate. However, replenishment experiments are hard to interpret simply in the synapse, because release can be limited either by the availability of vesicles or the availability of release sites, and in fact both priming and depriming rates affect pool sizes as well as recovery kinetics (Material and Methods, Eq. 13). Taking advantage of chromaffin cells (where priming sites are not limiting), we could show that Syt-7 has two effects: it decreases *k*_−*1*_ (the depriming rate) and it increases *k*_*1*_ (the priming rate) following stimulation (Fig. 4A-C; Table 1). In both cases the effect is approximately a factor of two. The physiological function is to upregulate the priming rate following stimulation to compensate for partial depletion of upstream vesicle pools. Recalculating the consequences of eliminating Syt-7 in a release site model (Fig. 4D-E), we showed that our findings are fully consistent with neuronal data showing overall slower recovery in Syt-7 KO neurons (Chen et al., 2017; Liu et al., 2014). The advantage of chromaffin cells is that we can derive priming and depriming rates separately, demonstrating that they are both affected by Syt-7.

### Syt-7 places dense-core vesicles at the plasma membrane

High-pressure-freezing 3D electron tomography in combination with classical 2D-EM showed that the total number of vesicles as well as the number of vesicles localized within 40 nm of the plasma membrane (i.e. membrane-proximal vesicles) were normal in the Syt-7 KO. Moreover, the number of membrane-attached vesicles (within 0-4 nm of the plasma membrane) was lower, although not significantly altered in Syt-7 KO chromaffin cells (Fig. 8P, T). In line with our previous findings for the analysis of Munc13-deficient chromaffin cells (Man et al., 2015), the calculated number of docked LDCVs per cell in the present study (~329 in WT cells) exceeds the number of vesicles in the functional RRP (~50 vesicles, according to Fig. 2A-B with 0.94 fF/vesicle (Pinheiro et al., 2014)) and SRP (~30 vesicles) at resting Ca^2+^-concentrations, thereby supporting the notion that functionally primed LDCVs in chromaffin cells are difficult to identify by morphological analysis due to a large number of dead-end docked LDCVs that are morphologically docked, but incapable of fusion upon stimulation (Hugo et al., 2013; Verhage and Sorensen, 2008). Indeed, it has been shown that by preventing full SNARE-zippering, synaptic vesicles can be rendered in a docked, but fusion-incompetent state (Gipson et al., 2017; Vardar et al., 2016). Interestingly, we found a significant reduction in the number of LDCVs within 0-6 nm of the plasma membrane in Syt-7 KO chromaffin cells. *In vitro* studies have shown that Munc13-1 forms a ~20 nm elongated structure that can bridge membranes (Quade et al., 2019; Xu et al., 2017) and individual SNARE-proteins can interact at distances smaller than 8 nm (Gao et al., 2012) or within 15 nm in the presence of SNARE-regulators like complexins (Li et al., 2011). The reduction of LDCVs in Syt-7 KO cells within 6 nm of the plasma membrane, which was accompanied by an increase in vesicle numbers within 20-40 nm, indicate a role for Syt-7 in mediating or stabilizing distance-dependent SNARE/Munc13-mediated interactions, thereby regulating vesicle priming and depriming. The increased number of more distant vesicles could account for the finding of a higher forward priming rate in Syt-7 KO when considering vesicles fusing 600 ms after stimulation (Table 1), and the increased size of the reserve pool in the hippocampal neurons of the Syt-7 KO (Duran et al., 2018).

A limitation of our studies is that Syt-7 and Syt-1 positive granules cannot be distinguished in EM tomograms, but a similar function for Syt-1 as a Ca^2+^-dependent distance-regulator between fusing membranes was previously proposed (Chang et al., 2018; van den Bogaart et al., 2011). In chromaffin cells, EM data obtained using chemical fixation methods implicated Syt-1 in vesicle docking (de Wit et al., 2009; Kedar et al., 2015). Although data obtained using chemical fixation cannot be compared directly to HPF-data [here and (Man et al., 2015)], overall these data indicate that either Syt-1 or Syt-7 might adopt the role of placing LDCVs into the vicinity of the plasma membrane to promote or stabilize SNARE/Munc13-interactions and thereby vesicle priming.

### Interdependence of Syt-7 and ubMunc13-2/phorbolester-dependent priming

We investigated how the priming role of Syt-7 interacts with ubMunc13-2, the dominating Munc13 protein in chromaffin cells (Man et al., 2015). Treatment of Syt-7 KO chromaffin cells with phorbolesters (which activate Munc13 proteins via their C1-domain) at low prestimulation [Ca^2+^] caused only a modest increase in RRP and SRP (overall release was potentiated by 21%), whereas phorbolesters caused a 126% increase in WT cells (Fig. 6). Likewise, in ubMunc13-2 overexpressing cells, the RRP was smaller in Syt-7 KO cells compared to WT cells (Fig. 7). These data show that the priming roles of Syt-7 and ubMunc13-2 are interdependent. To explain these data, as well as the EM-findings, we propose that a population of vesicles is attached to the plasma membrane via Syt-7. This brings the vesicles within the critical distance (6-10 nm) of the plasma membrane, where Munc13-proteins are able to bridge the gap between vesicle and plasma membrane, which allows SNARE-complex formation and priming (Fig. S11). In the Syt-7 KO, these vesicles reside at longer distances (20-40 nm), which prevents the formation of the SNARE-complex even when ubMunc13-2 is overexpressed. Interestingly, ubMunc13-2 overexpression stimulated a large SRP, but this SRP fused after a delay in almost all Syt-7 KO cells, whereas in Syt-7 WT cells - especially at higher prestimulation [Ca^2+^] - the SRP fused without delay. Thus, Syt-7 probably causes a movement towards the membrane, which is reinforced by binding to Ca^2+^ at < 1 μM; when Syt-7 is absent, vesicles reside at longer distances and fusion is therefore delayed. This can also explain the limited effect of treating Syt-7 KO cells with phorbolesters (Fig. 6), which anchor Munc13-2 into the plasma membrane. This will be ineffective if vesicles are out of reach of the Munc13-2/SNARE-complex. Finally, N-ethylmaleimide caused an increase in RRP size in Syt-7 KO cells when stimulated from a low prestimulation [Ca^2+^] (Fig. 5). This is consistent with our refilling experiments (Fig. 4) and indicates that a function of Syt-7 is to protect against depriming, either by binding directly to SNARE-proteins (Bacaj et al., 2015), or by moving vesicles closer to the membrane, where they can interact with Munc13-2, which in turn protects against depriming (He et al., 2017; Ma et al., 2013; Prinslow et al., 2019). The effect of NEM was only seen at low prestimulation [Ca^2+^], which indicates that other mechanisms stabilize the RRP at higher calcium concentrations. Overall, all three experiments (NEM, PMA and ubMunc13-2) are consistent with the misalignment of vesicles with the plasma membrane observed in the Syt-7 KO.

Although Ca^2+^-dependent priming *per se* is almost absent in the Syt-7 KO, the combination of Syt-7 KO with high prestimulation [Ca^2+^] and other manipulations to increase Munc13-priming (ubMunc13-2, PMA) allow priming in the Syt-7 KO, which is consistent with the presence of several priming pathways in chromaffin cells (Liu et al., 2010). If the main function of Syt-7 is to deliver vesicles to the membrane (and to Munc13) other calcium sensors that carry out similar functions will supplant the need for Syt-7. In chromaffin cells, Syt-1 (Nagy et al., 2006; Voets et al., 2001), Doc2B (Houy et al., 2017; Pinheiro et al., 2013) are the best candidates. As mentioned above, Syt-1 has been implicated in a similar priming role, also in neurons (Chang et al., 2018; Ruiter et al., 2019). Expression of membrane-bound Doc2B causes priming to saturate at the maximal level in chromaffin cells (Friedrich et al., 2008; Houy et al., 2017), i.e. the Ca^2+^-dependent priming process is overwhelmed, rendering priming Ca^2+^-independent, although Ca^2+^-dependent priming is still present in the Doc2B KO (Pinheiro et al., 2013).

We conclude that Syt-7 mediates Munc13- and Ca^2+^-dependent vesicle priming and vesicle fusion in a competitive and cooperative interplay with Syt-1, and that these functions involves positioning of vesicles in close membrane apposition.

## Materials and Methods

### Mouse lines and cell culture

Mouse lines (C57/Bl6-Syt-1, (Geppert et al., 1994), C57/Bl6-Syt-7 (Maximov et al., 2008)) were kept in an AAALAC-accredited stable at the University of Copenhagen operating a 12h/12h light/dark cycle with access to water and food ad libitum. Permission to keep and breed KO mice were obtained from the Danish Animal Experiments Inspectorate (permissions 2006/562-43 and 2018-15-0202-00157). Primary chromaffin cell cultures were prepared as described (Sorensen et al., 2003). Syt-7 knockout (KO) cells were obtained from P0-P2 pups of either sex originating from Syt-7 heterozygous crossing and identified by PCR genotyping (Maximov et al., 2008). Syt-1/Syt-7 DKO cells were obtained from embryos of either sex at embryonic day 18 (E18) by crossing mice that were homozygous KO (−/−) for Syt-7 and heterozygous (+/−) for Syt-1 (Schonn et al., 2008). The embryos were PCR genotyped for both Syt-1 and Syt-7. Cells marked Wild Type (WT) were P0-P2 WT littermates of the Syt-7 line, unless otherwise noted. The adrenal glands were dissected and cleaned in Locke’s solution consisting of (in mM) 154 NaCl, 5.6 KCl, 0.85 NaH_2_PO_4_, 2.15 Na_2_HPO_4_, and 10 glucose, and adjusted to pH 7.0. The glands were digested with 20-25 units/ml of papain enzyme for 45 min at 37 °C and 8% CO_2_ followed by 10-15 min inactivation with DMEM-Inactivation solution. To dissociate the cells, the glands were gently triturated and 40-50 μl of the cell suspension was placed onto the center of the coverslip, incubated to settle for 30 min and finally supplemented with 1-2 ml enriched DMEM media. Cells were used 2-5 days after plating. DMEM-Papain solution contained (in mM) (1 CaCl_2_, 0.5 EDTA) supplemented with 0.2 mg/ml l-cysteine and equilibrated with 8 % CO_2_. DMEM-Inactivation solution contained 10% heat-inactivated FCS, 2.5 mg/ml albumin, and 2.5 mg/ml trypsin inhibitor and equilibrated with 8% CO_2_. DMEM culture medium consisted of DMEM supplemented with 4 μl/ml penicillin/streptomycin and 10 μl/ml insulin-transferrin-selenium-X and equilibrated with 8% CO_2_.

### Viral constructs

The experiments made use of N-terminal pHluorin-tagged Syt-1 and Syt-7 (the α isoform (Fukuda et al., 2002)) constructs. The pHluorin (ecliptic EGFP) was preceded by a signal sequence of preprotachykinin to ensure correct orientation of the fusion protein into the vesicle membrane (Diril et al., 2006). The pHluorin-rat(rn)Syt cassette was cloned into the multiple cloning site of a lentiviral vector containing CMV promotor and a downstream WPRE sequence. To eliminate Syt-7 calcium binding, the C2 domains mutated fragments carrying aspartate/alanine exchange (C2A*: D225, 227, 233A; C2B*: D357, D359A) (Bacaj et al., 2013) were synthesised (Invitrogen GeneArt Gene Synthesis) and subcloned into the Syt-7 WT construct. In total, 5 constructs were produced: phluorin-rnSyt1; phluorin-rnSyt7α; phluorin-rnSyt7 α/C2A*; phluorin-rnSyt7 α/C2B*; phluorin-rnSyt7 α/C2AB*). All constructs were verified by sequencing. Lentiviruses were produced according to standard protocols using Lipofectamine2000 transfection and a HEK293FT cell host. The pHluorin-tags were used for identifying expressing cells. For lentiviral expression, cells were transduced 24 hours after prepping and incubated for 46-50 hours before being used for experiments. Acute expression, 4 – 6 hours, of EGFP-fused ubMunc13-2 was induced from a Semliki Forest Virus construct (Zikich et al., 2008).

### Immunostaining, confocal and structural illumination microscopy

Cells were plated on 25 mg/ml poly-D-lysine (Sigma P7405) coated coverslips. Prior to fixation, cells were cooled on ice for 3-5 min and fixated with ice cold 4% Paraformaldehyde (PFA; EMC 15710) for 15 min on ice, in the presence or absence of 0.2% Glutaraldehyde (Merck Millipore 104239), followed by 2% PFA for an additional 10 min at room temperature (RT). Cells were permeabilized with 0.15% Triton-X100 (Sigma-Aldrich T8787) for 15 min at RT and subsequently blocked with 0.2% cold fish gelatin (Sigma-Aldrich G7765), 1% goat serum (Thermo Fisher Scientific 16210064) and 3% Bovine Albumin Serum (Sigma-Aldrich A4503) for 1 h at RT. Cells were washed with PBS and glutaraldehyde autofluorescence was quenched with 0.1% Sodium Borohydride (NaBH4; Sigma-Aldrich 213462). Primary antibodies were diluted in blocking solution and incubated as follow: rabbit polyclonal α-Syt7 (1:500; Synaptic Systems SY105173) and mouse monoclonal α-Syt1 (1:500; Synaptic Systems SY105011) for 2 h at RT; rabbit polyclonal α-Syt1 (1:2000; W855; a gift from T. C. Südhof, Stanford, CA) and mouse monoclonal α-Syt7, clone 275/14 (1:200; Sigma-Aldrich MABN655) for 2 h at RT; rabbit polyclonal α-CgA (1:500; Abcam ab15160) and mouse monoclonal α-Syt7, clone 275/14 (1:200; Sigma-Aldrich MABN655) or mouse monoclonal α-Syt1 (1:500; Synaptic Systems SY105011), overnight at 4°C. Secondary antibodies used were goat α-chicken Alexa Fluor 488 conjugate (1:500; Abcam ab150169), goat α-mouse Alexa Fluor 488 conjugate (1:500, Thermo Fisher Scientific A11029), goat α-mouse Alexa Fluor 546 conjugate (1:500, Thermo Fisher Scientific A11003) and goat α-rabbit Alexa Fluor 647 conjugate (1:500; Thermo Fisher Scientific A21245). Immunofluorescence was visualized using a Zeiss LSM 780 inverted confocal with oil-immersion Plan-Apochromat NA 1.4 63x objective. The fluorophores were excited with Argon 488 nm (25 mW), HeNe 543 nm (1.2 mW), 633 nm (5 mW), and intune 488-645 nm (5 mW) lasers. Linear unmixing was applied on cells stained with more than two fluorophores in Zen Black Zeiss software. Control cells stained with a single fluorophore were used to define the spectral finger print. Quantification of Syt-1, Syt-7 was performed on ImageJ software on average projections of 0.5 μm increment z-stacks. Mean intensity of circular cell ROI was background subtracted and the value was normalized to control cells (wild type cells or Syt7-KO cells). Control cells were acquired on the same day as sample cells, and laser power, gain and emission detection were unchanged. Estimation of colocalization of Syt-1, Syt-7 and CgA was possible by applying the Mander’s Coefficient on single z-stacks, using the JACoP plugin for ImageJ.

For further colocalization analysis (Fig. S10), Syt-7 WT cells immunostained against Syt-1 and Syt-7 were imaged with 1 μm Z-stack increment. Single vesicles in each slice were selected in the Syt-1 staining by automatically finding a maxima and drawing a square ROI around it in a custom-written ImageJ macro. The selected ROIs were averaged; to gain a line plot the 3 middle lanes of the image were averaged. The identified ROIs were then applied to the Syt-7 staining channel to create a corresponding plot for Syt-7. On average, 299 vesicles were selected and averaged per cell.

For 3D-Structural Illumination Microscopy (3D-SIM), images with voxel size (x,y,z) in μm: 0.03 x 0.03 x 0.11) were obtained with a Zeiss Elyra PS.1 microscope equipped with a sCMOS PCO.edge camera and an oil-immersion Plan-Apochromat NA 1.4 63x objective. Alexa Fluor −488 and −647 were exited with 488 HR Diode-200mW and HR Diode-150mW lasers, respectively. Distance-based co-localization analysis was performed using Spots colocalization (ComDet) ImageJ plugin (available online, authored by Eugene Katrukha). Vesicles were automatically detected in their respective channels (Syt-1 and Syt-7) by setting the detection threshold to 3xSD above background noise and an approximate particle size of 120 nm. The approximate particle size was based on the FWHM of a Gaussian fit of averaged vesicles where the averaging routine described above was applied (FWHM of 1200 averaged Syt-1 vesicles: 130 nm; FWHM of 600 averaged Syt-7 vesicles: 149 nm). Note that the 120 nm particle size setting allows detection of vesicles that are larger than 120 nm in diameter but restrict the detection of very large undefined structures. Finally, vesicles were scored as co-localized if the coordinates from one channel were within 60 nm distance (half of the set particle size) in the other channel. Since the vesicles are larger than one imaging plane (the vesicle diameter is ~175 nm and would be increased further by antibody binding), they would show up in more than one imaging plane, and we therefore analyzed every third plane, equivalent to 330 nm Z-intervals. To estimate the number of duplicate vesicles (same vesicle that is detected in two subsequent planes) we compared the XY coordinates of vesicles in sequentially analyzed planes and counted matching positions as duplicates. This indicated that the number of duplicates were <1%.

### Western blot

Adrenal glands and brain extracts were collected from P0-1 Syt-7 WT and KO mice and lysed in RIPA buffer supplemented with Protease Inhibitor Cocktail (Invitrogen, 89900). The supernatants were collected and protein concentrations were estimated by using the BCA Protein Assay Kit (Pierce 23227) and plotting the resulting BSA curve. 25 μg and 15 μg of protein, from adrenal glands and brain extracts, respectively, were resolved by 4-12% SDS-PAGE (Invitrogen, Thermo Fisher Scientific) and wet-transferred onto an Amersham Hybond LFP PVDF membrane (GE Healthcare). The membrane was blotted with rabbit polyclonal α-Syt7 (1:500; Synaptic Systems SY105173) and mouse monoclonal α-VCP (1:10000; Abcam ab11433), as a loading control, followed by α-rabbit (1:2000; Invitrogen P0448) and HRP-conjugated α-mouse (1:2000; Invitrogen P0447) secondary antibodies. The blot was developed by chemiluminescence SuperSignal™ West Femto and Pierce™ ECL Plus Western blotting substrate systems (Thermofisher Scientific) and immunoreactive bands were detected using the FluorChemE image acquisition system (ProteinSimple) equipped with a cooled CCD camera.

### Electrophysiology

Exocytosis was monitored by combining membrane capacitance measurements and carbon fibre amperometry. Capacitance measurements were based on the Lindau-Neher technique using Pulse HEKA software with Lock-In extension. A 70 mV peak-to-peak sinusoid (1000 Hz) was applied around a holding potential of −70 mV in the whole-cell configuration. The clamp currents were filtered at 3 kHz and recorded at 12 kHz with an EPC9 HEKA amplifier. Secretion was triggered by 1-2 ms UV flash-photolysis of the caged Ca^2+^ compound nitrophenyl-EGTA, infused through the patch pipette. The UV-flash delivered from a flash lamp (Rapp Optoelectronic, JML-C2) was bandpass-filtered around 395 nm, transmitted through a light guide and a dual condenser and focused with a Fluar 40X/N.A. 1.30 oil objective.

The intracellular Ca^2+^ concentration was determined as described in (Nagy et al., 2002). Two florescent dyes with different affinities toward Ca^2+^, Fura4F (Kd=1 μM) and furaptra (Kd=40 μM) were infused via the pipette into the cell. For ratiometric detection, alternating monochromator excitations of 350 nm and 380 nm were generated at 40 Hz and emission was detected via a photodiode, recorded at 3 kHz and filtered at 12 kHz. The 350/380 ratio was pre-calibrated by infusing the cell with known Ca^2+^ concentrations.

Amperometric recordings were performed as previously described (Bruns, 2004) using a carbon fibre (5-10 μm diameter) insulated with polyethylene and mounted in glass pipette. The fibre was clamped at 700 mV, currents were filtered at 5 kHz and sampled at 25 kHz by an EPC7 HEKA amplifier. 50 Hz noise was online eliminated by a Humbug noise eliminator device. For single spike analysis, amperometric traces were off-line filtered at 500 Hz using a Gaussian filter, threshold detected at 5 pA and analysed with a Igor Pro macro (Mosharov, 2008). The interspike interval is the median time interval between the spikes (Mosharov, 2008).

The pipette solution contained (in mM): 100 Cs-glutamate, 8 NaCl, 4 CaCl_2_, 32 Cs-HEPES, 2 Mg-ATP, 0.3 GTP, 5 NPE, 0.4 fura-4F, 0.4 furaptra, and 1 vitamin C. Adjusted to pH 7.2 and osmolarity to ~295 mOsm. The extracellular solution contained (in mM): 145 NaCl, 2.8 KCl, 2 CaCl_2_, 1 MgCl_2_, 10 HEPES, and 11 glucose. Adjusted to pH 7.2 and osmolarity to ~305 mOsm. For the double flash (recovery) experiment (Fig. 4 and Fig. S4), the NPE concentration was reduced to 3 mM and UV flash intensity was adjusted before the second flash to ensure comparable post-flash calcium levels and fast Ca^2+^ relaxation after the flash. *N*-Ethylmaleimide (NEM) (Sigma 04259) was prepared fresh prior to each experiment and added in the pipette solution to a final concentration of 200 μM. NEM was infused into the cells for 60 – 100 s before stimulation. Control cells were treated equally but patched with a pipette solution that did not contain NEM. Phorbol 12-myristate 13-acetate (PMA) (Sigma P8139) was dissolved in DMSO and diluted in extracellular solution immediately prior to the experiment, to a final concentration of 100 nM and used within 2 hours.

### Kinetics analysis

Pool sizes were determined either 0.5 s after the flash and designated as “Burst” or by fitting the capacitance trace with a sum of two exponentials plus a straight line using a custom written Igor Pro macro (Wavemetrics):

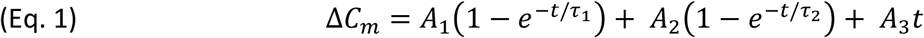

where the amplitudes A_1_ and A_2_ are the sizes of the releasable pools, and τ_1_ and τ_2_ are their fusion time constants. The resulting exponential components were assigned as RRP or SRP according to the estimated time constant (τ). Except if noted otherwise, kinetic components were considered to originate from the RRP when τ ≤ 60 ms and from the SRP when 60 ms ≤τ ≤ 1000 ms. The sustained release rates were calculated as the linear rate (*A*_*3*_) following fusion of SRP and RRP. In cases where the fit identified two time constants within the same cut off criterion (i.e. both time constants would correspond to either a RRP or the SRP), the trace was refitted with a single exponential for the corresponding component.

In one case (Munc13-2 overexpression in Syt-7 KO and sometimes in Syt-7 WT), capacitance traces had an S-formed shape, which made it impossible to fit them with (Eq.1). Instead, we here derive a new function for the SRP, which takes into account the delayed fusion of this pool of vesicles. We first observe (in agreement with Eq. 1) that the fusion of the SRP follows the evolution:

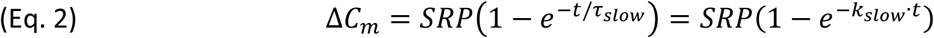

where *τ*_*slow*_ is the time constant of fusion and *k*_*slow*_ is the rate constant of fusion and 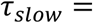 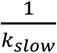. Let us assume that the fusion rate of SRP is not instantaneously at its max value, *k*_*slow*_; instead, the fusion rate of the SRP, *k*_2_, increases gradually towards *k*_*slow*_:

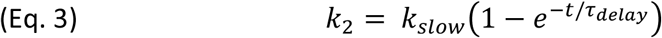

where *τ*_*delay*_ is the time constant of the development of *k*_2_ towards the value *k*_*slow*_. Therefore, the new model for fusion of the SRP is:

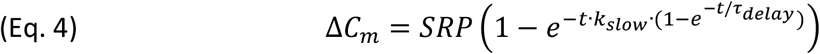

Fitting this equation to capacitance traces is an illformed problem, because *k*_*slow*_ and *k*_*delay*_ can not both be determined. To see why, we consider the Taylor expansion of the expression for *k*_2_:

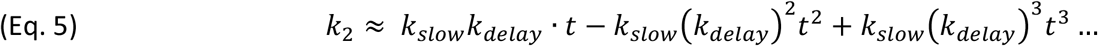

If the delay is long (i.e. *k*_*delay*_ is small), we can replace *k*_2_ with its first-order approximation, *k*_*slow*_*k*_*delay*_ · *t* to yield

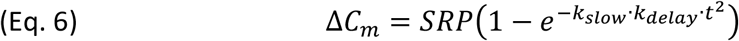

Or, if we set 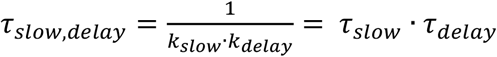 we get

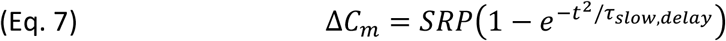

Here, *τ*_*slow,delay*_ has a different meaning than *τ*_*slow*_, because it includes both the delay and the fusion kinetics of the SRP. For fitting with a delayed SRP, Eq. 7 was substituted into Eq. 1.

### Priming models

#### Model I – no maximal primed vesicle pool size

To understand the consequences of changes to priming and depriming rates, we considered a simple 2-pool model, similar to (Heinemann et al., 1993), where a large reserve pool (‘DP’ for Depot Pool) is filling up a primed vesicle pool, (‘PP’ for Primed Pool), through a reversible priming step, which drives priming forward with rate constant *k*_*1*_, and supports depriming (i.e. the reverse priming reaction) with rate constant *k*_−*1*_. Fusion is supported with rate constant *k*_*f*_. Under such conditions, the change in the primed pool is given by:

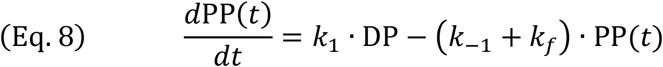

If we assume that the DP does not change size during the experiment, we get the general solution:

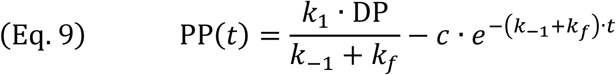

where *c* is an arbitrary constant.

If we insert the initial condition PP(0) = 0, we get the specific solution relevant for a pool recovery experiment:

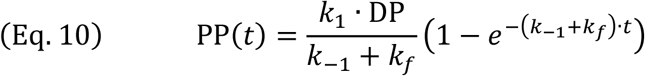

where the PP size at equilibrium (t = ∞) is

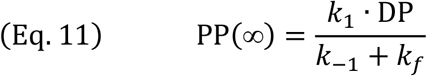

#### Model II – with a maximal primed pool size (release site model)

If we assume that the Primed Pool is limited by the number of release sites, we can modify (Eq. 8) to include a maximal pool size, *PP*_*max*_ (a constant). If we further assume that those release sites that are vacated by fusion, or depriming, are immediately available again for priming, the model becomes

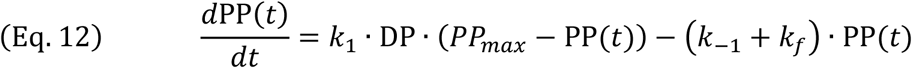

In reality, the release sites will take a while to recover after fusion (Hosoi et al., 2009); however, we shall consider the simpler situation. Note that *k*_*1*_ has now changed unit from *s^−1^* to *release site^−1^ s^−1^* (or *fF^−1^s^−1^*, in capacitance units).

For a pool recovery experiment (initial condition PP(0) = 0), we get the solution

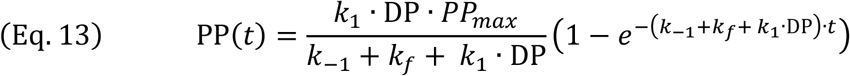

where the time constant for recovery (1/(*k*_−1_ + *k*_*f*_ + *k*_1_ · DP)) now is sped up by the forward priming rate (this is a difference to *Model I*). The steady-state solution is

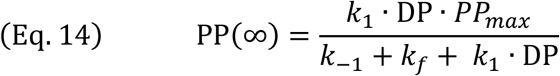

#### Comparison of Model I and II (Figure 4 and S3)

We identified the depriming rate (*k*_−*1*_) for WT and Syt-7 KO chromaffin cells by fitting a normalized version of (Eq. 10) to a recovery curve, under the assumption that *k*_*f*_ = 0 (Fig. 4B-C). When considering recovery at 60 ms after stimulation (corresponding approximately to the RRP), the results showed that *k*_−*1*_ was increased by a factor of 2.12 in the Syt-7 KO (from 0.043 s^−1^ in the WT to 0.091 s^−1^ in the KO, Fig. 4B). Comparison to the pool size obtained by the first stimulation then made it possible to calculate the forward priming rate before stimulation, *k*_1_ · DP (from Eq. 11), which was almost unchanged (3.21 fF/s in the WT and 3.07 fF/s in the Syt-7 KO). To take into account the slight overfilling during recovery in the WT we assumed that this would result from a change in priming after stimulation. To calculate the *k*_1_ · DP after the stimulation, we multiplied *k*_1_ · DP by the fitted normalized Plateau value (yielding 3.78 fF/s in WT and 1.90 fF/s in the Syt-7 KO). The recovery data at 600 ms were treated in the same way to identify *k*_−*1*_, and *k*_1_ · DP before and after stimulation. All values are given in Table 1.

In order to understand how the presence of release sites would change the observation in the Syt-7 KO, we searched for a solution to Model II with unchanged recovery time constant and pool size in the WT case. For recovery assessed at 60 ms after stimulation, we first assumed that the RRP is at 90% capacity at rest (*k*_*f*_ = 0), which identifies *PP*_*max*_ = 74.2 fF/0.9 = 82.4 fF. We now isolated *k*_1_ · DP and *k*_−*1*_ from the two equations (where we again assume *k*_*f*_ = 0 during recovery):

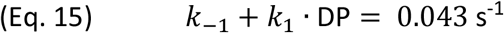

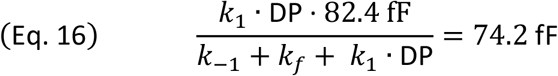

which yields *k*_1_ · DP = 0.039 s^−1^ and k_−1_ = 0.0043 s^−1^. Simulating a recovery experiment with these parameters in Model II yielded a curve identical to the one in Model I with WT parameters (Fig. 4E). To understand the consequences of changing the depriming and priming rates, we increased *k*_−*1*_ by a factor of 2.12 and fixed pre- and post-stimulation *k*_1_ · DP at the values found in the Syt-7 KO (Table 1). This yielded the calculated recovery curves for the Syt-7 KO in a cell with release sites (Fig. 4E). Predicted recovery at 600 ms in a release site model was calculated in the same way (Fig. 4F).

### Electron Microscopy

Samples for the ultrastructural analysis of LDCV docking in chromaffin cells were prepared according to a published protocol (Man et al., 2015) with only minor modifications. Briefly, adrenal glands were removed from Syt-7 KO and WT littermate P0 pups and sectioned into 100μm-thick slices using a vibratome. Adrenal gland slices were allowed to recover for 15 min at 37°C in bicarbonate-buffered saline in the presence of 0.2 mM (+)-tubocurarine and then kept at room temperature in the same solution before cryofixation in external cryoprotectant (20% bovine serum albumin in bicarbonate-buffered saline) using a HPM100 HPF device (Leica). Freeze substitution was performed as previously published (Rostaing et al., 2006) and samples were embedded in EPON resin for 24 hours at 60°C. Ultrathin (60 nm, for 2D analysis) and semithin (350 nm, for 3D analysis) sections were collected onto Formvar-filmed, carbon-coated copper mesh grids. Ultrathin sections were poststained with uranyl acetate and lead citrate before imaging. Semithin sections were briefly incubated in Protein A conjugated to 15 nm gold particles (Cell Microscopy Center, Utrecht, The Netherlands). 2D- and 3D-EM imaging and the analysis of LDCV docking in chromaffin cells was performed exactly as previously published (Man et al., 2015).

### Calculation of total number and number of docked LDCVs in chromaffin cells

To calculate the total number of LDCVs per WT cell, we used the 2D analysis, together with the mean diameter of the vesicles, as determined from 3D-analysis (175.5 ± 4.1 nm). We use the cytoplasm density of vesicles (*δ*_v_ = 4.71 ± 0.18 vesicles/μm^2^) and proceed to calculate the volume fraction of LDCVs in the cytoplasm. Each ultrathin section is 60 nm thick (*h*). We first assume that each vesicle profile within a slice occupy a cylinder-shape with a height of 60 nm and a diameter of 0.1755 μm (vesicular radius *r*_*ves*_ = 0.08775 μm). The cylinder-shape occupies a volume of π x (0.08775μm)^2^ x 0.060 μm = 0.00145 μm^3^. The total volume of a 1 μm^2^ area of the slice is 0.060 μm^3^, and the (uncorrected) volume-fraction of LDCVs would be 4.71 x 0.00145 μm^3^ / 0.060 μm^3^ = 0.114. However, we need to correct for the spherical shape and tangential slicing of vesicles, which will lower the volume-fraction. The volume of a LDCV is (4/3) x π x (0.08775 μm)^3^ = 0.002830 μm^3^. The volume of the circumscribed cylinder is π x (0.08775μm)^2^ x (0.1755+0.060) μm = 0.00570 μm^3^, where we have assumed that a LDCV is identified as such when 30 nm of a 60 nm section cuts the vesicle tangentially, and thus we add 2 x 30 nm = 60 nm to the cylinder height. Since cutting the vesicle at any point is equally likely, we apply the volume correction 0.00283 μm^3^/0.00570 μm^3^ = 0.497. Thus, the corrected volume-fraction of LDCVs is 0.114 x 0.497 = 0.0567. The radius of a chromaffin cell was determined from the mean cell capacitance measured in patch-clamp experiments (3.83 ± 0.057 pF). With a specific capacitance of 10^−2^ F/m^2^, we get a plasma membrane (PM) area of 383 μm^2^. Thus, the radius of a chromaffin cell is 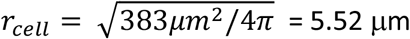. We assume that the nucleus’ radius is half of the cell radius, i.e. 2.76 μm, which is comparable to the radius determined from the nucleus area in 2D EM micrographs (2.63 ± 0.06 μm), although the latter value will be affected by tangential slicing of the nucleus. The volume of the cytoplasm is then (4/3) π (5.52 μm)^3^ - (4/3) π (2.76 μm)^3^ = 704.5 μm^3^ – 88.1 μm^3^ = 616.4 μm^3^. The total volume of LCDVs is therefore 0.0567 x 616.4 μm^3^ = 34.95 μm^3^, which yields 34.95 μm^3^ / 0.002830 μm^3^ = <u>12,334 LDCVs per Syt-7 WT cell</u>. When assembling the considerations above into a single equation, the number of vesicles per cell (*N*_*v*_) is given by

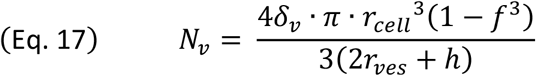

where *δ*_v_ is the density of vesicles per μm^2^ cytoplasm, *r*_*cell*_ is the radius of the cell, *r*_*ves*_ is the radius of a vesicle, *h* is the thickness of an ultrathin section and *f* is the ratio of nucleus to cell radius (here we used *f* = 0.5). The corresponding numbers for Syt-7 KO are: LDCV diameter = 165.3 ± 3.4 nm, cytoplasm density of LDCVs = 4.70 ± 0.23 vesicles/μm^2^, resting cell capacitance = 3.91 ± 0.062 fF, which yields a total of <u>13,270 LDCVs per Syt-7 KO cell</u>.

The number of membrane-proximal LDCVs per cell (*n*_*a*_, defined as vesicles within 40 nm of the cell membrane) can be calculated from the number of LDCVs per μm plasma membrane length (*n*_*l*_) using the formula *n*_*a*_ = *n*_*l*_/(*d*_*v*_ + 0.06), where *d*_*v*_ is the vesicle diameter (Parsons et al., 1995; Plattner et al., 1997). Our 3D-ET approach yields accurate estimates of *d*_*v*_ (see above). For the Syt-7 WT, we estimated *n*_*l*_ = 0.75 ± 0.05 vesicles/μm, which yields *n*_*a*_= 3.18 vesicles per μm^2^. With a PM area of 383 μm^2^ (see above), we get a total of ~1219 membrane-proximal (<40 nm) vesicles per cell. From 3D-ET, we know that 27 ± 2.5 % of vesicles within 40 nm of the PM are docked (i.e. physically attached to the plasma membrane), which means <u>Syt-7 WT cells have ~ 329 docked vesicles/cell</u>. For the Syt-7 KO we have estimated *n*_*l*_ = 0.73 ± 0.05 vesicles/μm, which gives us *n*_*a*_= 3.24 vesicles per μm^2^. With a PM area of 391 μm^2^ we have a total of ~1267 vesicles within 40 nm of the PM. From 3D-ET 21±2.5% are membrane-attached, which means that <u>Syt-7 KO cells have ~ 266 docked vesicles/cell</u>.

### Statistics

The data are presented as mean ± SEM; N indicates the number of cells. Non-parametric Mann-Whitney or Kruskal-Wallis with post Dunn’s test were applied for all capacitance measurements and release time constants. For other data, student’s *t*-test or One-way ANOVA with post-hoc Tukey’s test or Dunnett’s test were applied on data with similar variances. The variances were analyzed with F test for two-sample data or with Bartlett’s test for comparing more than two data samples. Heteroscedastic data were log transformed to satisfy the prerequisite of homogeneous variances. Non-parametric Mann-Whitney or Kruskal-Wallis with post Dunn’s test were applied on data that failed to meet the criteria for parametric test after log-transformation.

### Online supplementary material

Fig. S1 quantifies the total amperometric charge measured in each Ca^2+^-uncaging experiments. Fig. S2 demonstrates overexpression of Syt-7 WT protein, and Syt-7 with mutated Ca^2+^-binding sites in the C2A (C2A*), C2B (C2B*) or both the C2A and C2B domain (C2AB*). Mutation of either or both C2-domains eliminates the effect of Syt-7 on priming. Fig. S3 shows how changes in priming and depriming rates affects recovery of the primed vesicle pool in the absence and presence of release sites. Fig. S4 shows data from double stimulation experiments in Syt-7 WT and Syt-7 KO cells. Fig. S5 shows the effect of applying phorbolester to Syt-7 WT and KO cells at high prestimulation [Ca^2+^]. Fig. S6 shows the effect overexpressing ubMunc13-2 in Syt-7 WT and KO cells at higher prestimulation [Ca^2+^]. Fig. S7 show that there is no effect of NEM when Syt-7 WT or KO are stimulated from a higher prestimulation [Ca^2+^]. Fig. S8 shows double labeling of Syt-1 and chromogranin and Syt-7 and chromogranin, respectively. Fig. S9 shows controls for double staining of Syt-1 and Syt-7, staining with another antibody combination, and Western blot of Syt-7 WT and KO adrenal glands and brain extracts. Fig. S10 demonstrates that averaging confocal images around Syt-1 positive vesicular structures reveals a colocalized Syt-7 signal. Fig. S11 illustrates the proposed model for Syt-7 and ubMunc13-2 dependent vesicle priming.

## Acknowledgements

We would like to thank Anne Marie Nordvig Petersen and Dorte Lauritsen for expert technical assistance. This investigation was supported by the University of Copenhagen 2016 (KU2016) excellence program, the Novo Nordic Foundation, the Danish Medical Research Council (all JBS), and the Lundbeck Foundation (JBS and PSP). We thank Erwin Neher for commenting on an earlier version of the manuscript.

