## Supplemental Figures for "Synaptotagmin-7 places dense-core vesicles at the cell membrane to promote Munc13-2- and Ca^2+^-dependent priming"

[4] Present address: Department of Neuroscience, University of Copenhagen, 2200 Copenhagen N, Denmark.

[5] Equal contribution.

**Content:**

Figure S1-S11.

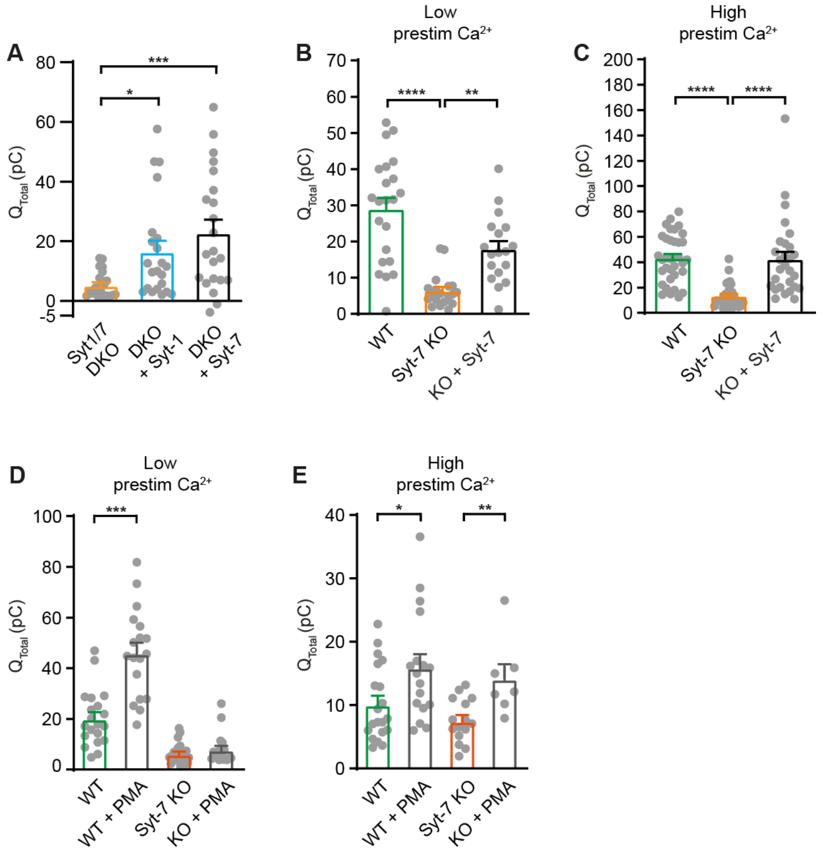


**Figure S1 – Amperometric charge quantification**

**A** Supplemental to Figure 1: Integrated amperometric current 5 s after $Ca^{2+}$-uncaging (mean ± SEM) of Syt-1/Syt-7 double KO (DKO), and DKO cells overexpressing Syt-1 (DKO + Syt-1) or Syt-7 (DKO + Syt-7). *: p<0.05; ***: p<0.001. Kruskal-Wallis test with post-hoc Dunn's test.

**B** Supplemental to Figure 2A: Integrated amperometry (mean ± SEM) of WT, Syt-7 KO and Syt-7 KO overexpressing Syt-7 (KO + Syt-7) stimulated from low prestimulation [$Ca^{2+}$]. **: p<0.01; ****: p<0.0001.Kruskal-Wallis test with post-hoc Dunn's test.

**C** Supplemental to Figure 2F: Integrated amperometry (mean ± SEM) of WT, Syt-7 KO and KO + Syt-7 stimulated from high prestimulation [$Ca^{2+}$]. ****: p<0.0001. Kruskal-Wallis test with post-hoc Dunn's test.

**D** Supplemental to Figure 6: Integrated amperometry (mean ± SEM) of WT, WT cells treated with 100 nM phorbol 12-myristate 13-acetate (PMA) (WT + PMA), Syt-7 KO and Syt-7 KO with 100 nM PMA (Syt-7 KO + PMA) stimulated from low prestimulation [$Ca^{2+}$]. ***: p<0.001. Student's *t*-test.

**E** Supplemental to Figure S5: Integrated amperometry (mean ± SEM) of WT, WT cells perfused with 100 nM phorbol 12-myristate 13-acetate (PMA) (WT + PMA), Syt-7 KO and Syt-7 KO treated with 100 nM PMA (syt-7 KO + PMA) stimulated from high prestimulation [$Ca^{2+}$]. *: $p<0.05$; **: $p<0.01$. Student's *t*-test.

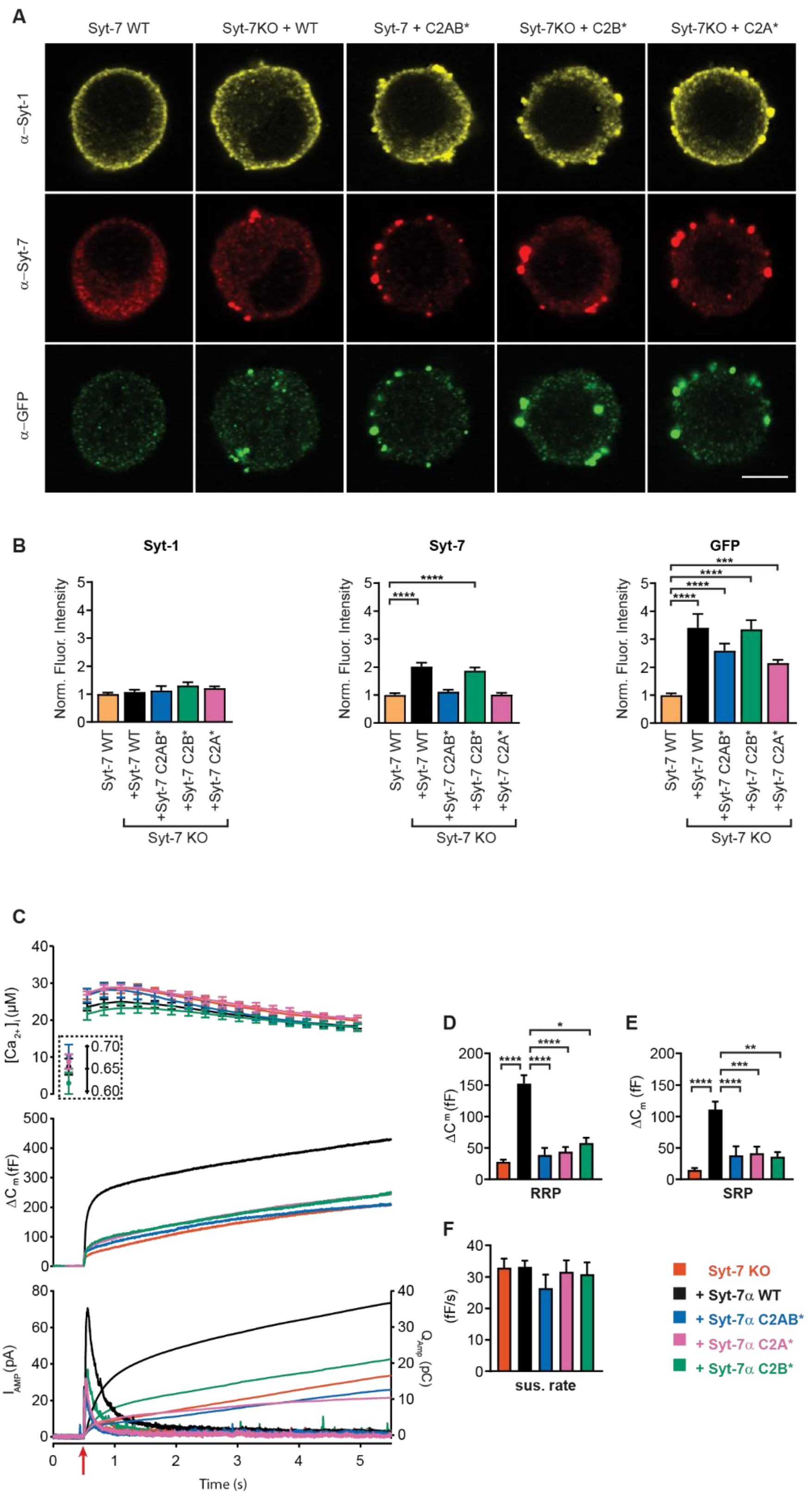


**Figure S2 – Mutation of $Ca^{2+}$-binding sites in Syt-7 abolishes rescue function.**

**A** Single confocal slices of new-born mouse chromaffin cells stained against Syt-1 (α-Syt-1), Syt-7 (α-Syt-7) and GFP (α-GFP) in WT cells, Syt-7 KO cells and Syt-7 KO cells overexpressing Syt-7 WT, a Syt-7 C2A-mutation (C2A*)(D225,227,233A), a Syt-7 C2B-mutation (C2B*) (D357,D359A), and a Syt-7 mutated in both C2A and C2B (C2AB*) (D225,227,233,357,359A)

**B** Quantification (mean ± SEM) of staining against Syt-1, Syt-7 and GFP normalized to Syt-7 WT cells. ***: p<0.001; ****: p<0.0001. In B, middle panel: One-way ANOVA with post-hoc Dunnett's test; B, left and right panels: Kruskal-Wallis test with post-hoc Dunn's test). Number of cells: Syt-7 WT: N = 20 cells; Syt-7 KO overexpressing Syt-7: N= 21; C2A* mutation: N = 20 cells; C2B* mutation: N = 20 cells; C2AB* mutation: N = 20 cells.

**C** Calcium uncaging experiment from a relatively high prestimulation [$Ca^{2+}$] in Syt-7 KO cells (vermilion), in Syt-7 KO cells overexpressing Syt-7 WT (black traces), a Syt-7 C2A-mutation (C2A*, pink traces, D225,227,233A), a Syt-7 C2B-mutation (C2B*, green traces D357,D359A), and a Syt-7 mutated in both C2A and C2B (C2AB*, blue traces, D225,227,233,357,359A). Panels are arranged as in Fig. 1A.

**D** Sizes (mean ± SEM) of the RRP.

**E** Sizes (mean ± SEM) of the SRP.

**F** Sustained rate (mean ± SEM) of secretion.

*: p<0.05; **: p<0.01; ***: p<0.001; ****: p<0.0001. Kruskal-Wallis with post-hoc Dunn's Multiple Comparison test. Syt-7 KO: N = 50 cells; Syt-7 KO + Syt-7 WT: N = 52 cells, Syt-7 KO + Syt-7 C2A*: N = 30 cells, Syt-7 KO + Syt-7 C2B*: N = 22 cells, Syt-7 KO + Syt-7 C2AB*: 20 cells.

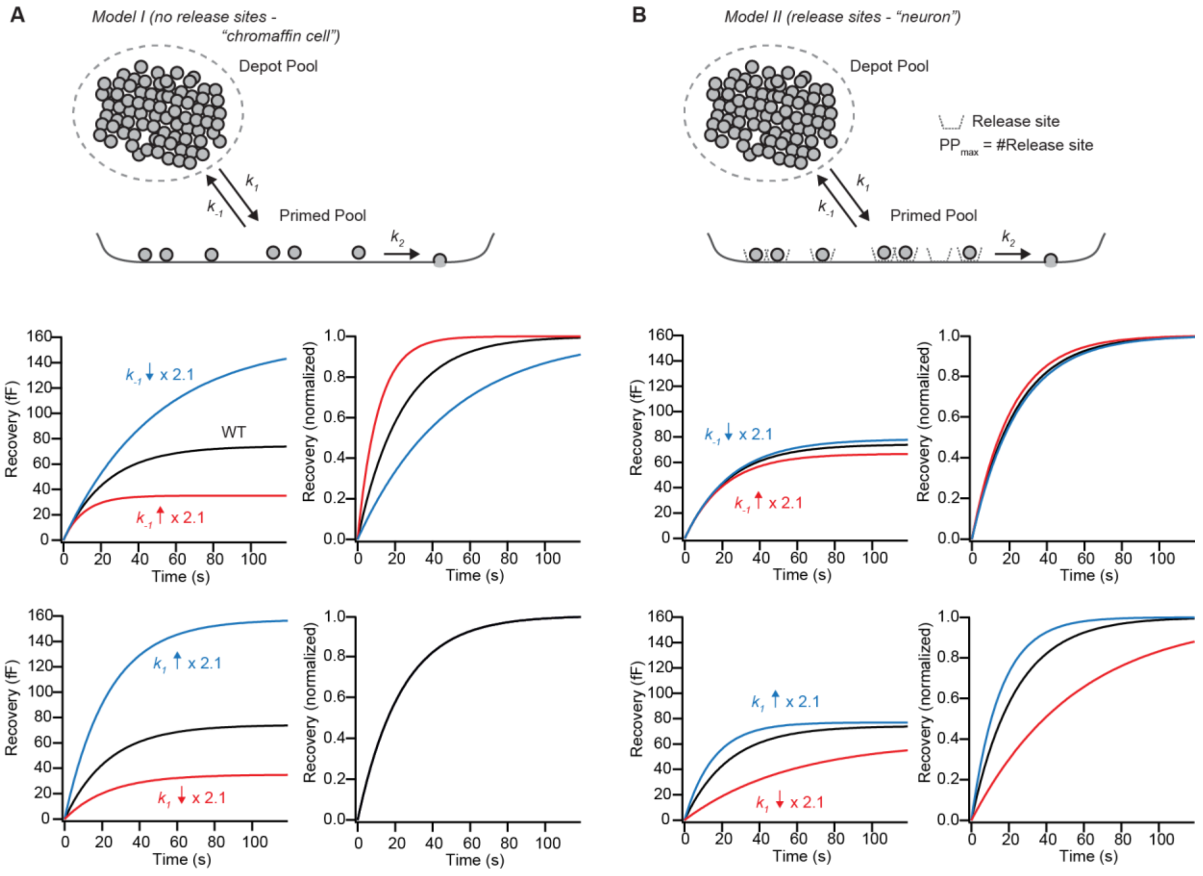


**Figure S3 – Differences in unpriming rate can be distinguished during pool recovery, if the Primed Pool is not limited by release sites**

**A** In *Model I*, we assume that vesicles from a very large Depot Pool prime reversibly (priming rate: $k_1$; unpriming rate: $k_{-1}$) into the Primed Pool, which is free to change is size. Fusion happens from the primed pool with rate $k_f$. The bottom left panel shows the recovery of the primed pool (black line) with parameters from a fit to WT data (Fig. 4B), $k1 \cdot$ DP = 3.21 fF/s, $k_{-1}$ = 0.043 $s^{-1}$, and after increasing (red line, $k_{-1}$ = 0.091 $s^{-1}$) or decreasing (blue line, $k_{-1}$ = 0.020 $s^{-1}$) the $k_{-1}$ by a factor of 2.12. The red line corresponds closely to the fit of the Syt-7 KO condition. Right-hand panel: after normalization, it is appreciated that the high-$k_{-1}$ condition recovers with faster kinetics than the WT condition, which recovers faster than the low-$k_1$ condition.

**B** In *Model II*, we assume that the Primed Pool is limited by a fixed number of release sites, which sets an upper limit ($PP_{max}$) to the Primed Pool size. The bottom panel shows the recovery of the primed pool (black line) with parameters fitting the Syt-7 WT condition (black curve, see Materials and Methods), and after increasing (red curve) or decreasing (blue curve) the unpriming rate ($k_{-1}$) by a factor of 2.1. Right-hand panel: normalized traces. In the presence of release sites, recovery becomes less sensitive to $k_{-1}$.

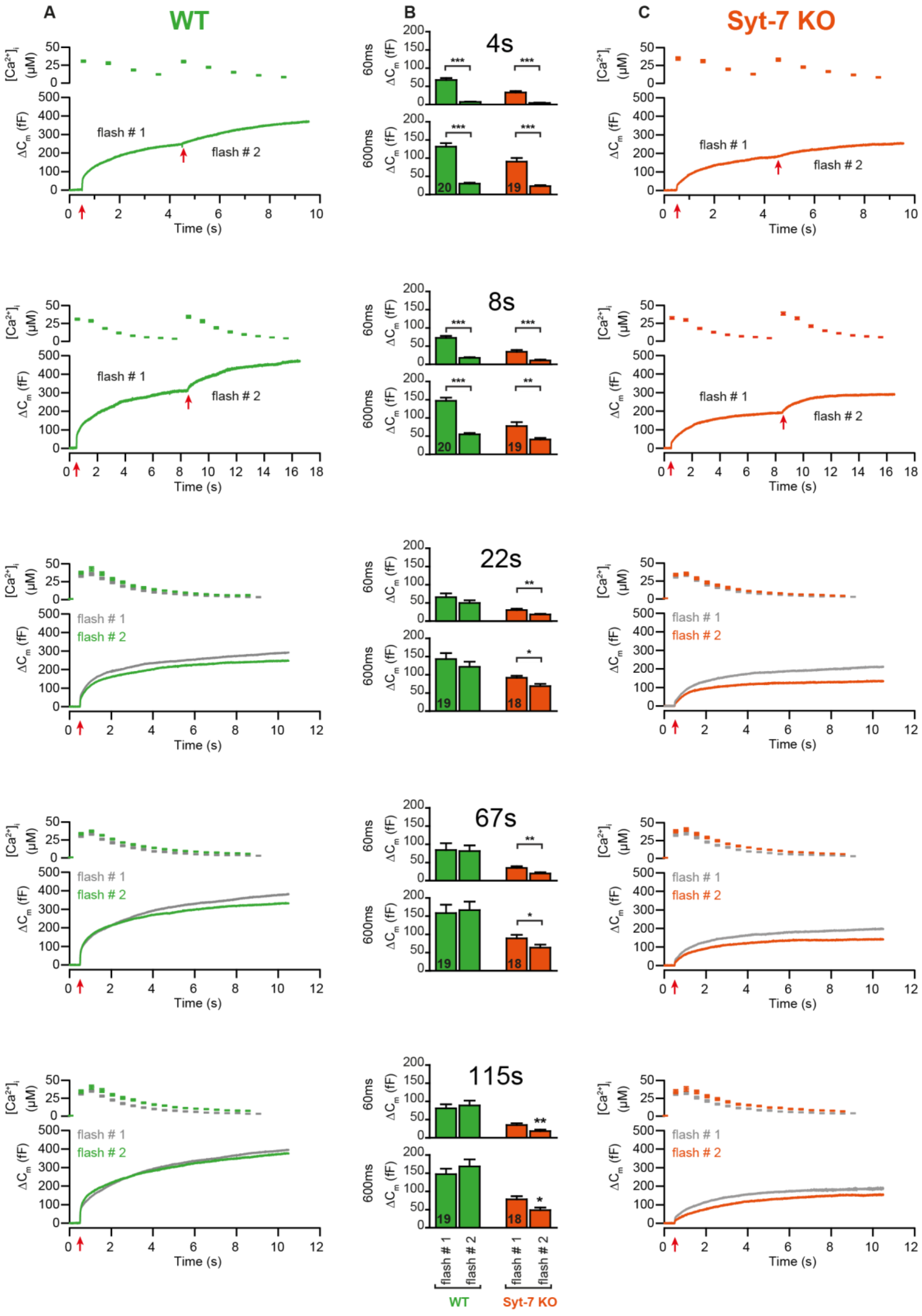

A
WT
B
4s
8s
22s
67s
115s
C
Syt-7 KO
flash # 1
flash # 2
Time (s)
[Ca2+]i (µM)
ΔCm (fF)
60ms
600ms
WT
Syt-7 KO

**Figure S4 – Double stimulation experiments in Syt-7 WT and KO cells.**

**A** Mean $[Ca^{2+}]_i$ and capacitance traces of Syt-7 WT cells during double uncaging experiments. Time intervals are indicated in panel B. For time intervals 4 and 8 s, the two sequential stimulations are shown on a continuous time axis, for longer intervals they are shown overlaid.

**B** Quantification of capacitance at 60 and 600 ms after flash uncaging, for the first and second stimulation. Number of cells (N) are shown on bar diagrams

**C** Mean $[Ca^{2+}]_i$ and capacitance traces of Syt-7 KO cells during double uncaging experiments. Arranged as in A. *: $p<0.05$; **: $p<0.01$; ***: $p<0.001$. Mann-Whitney test.

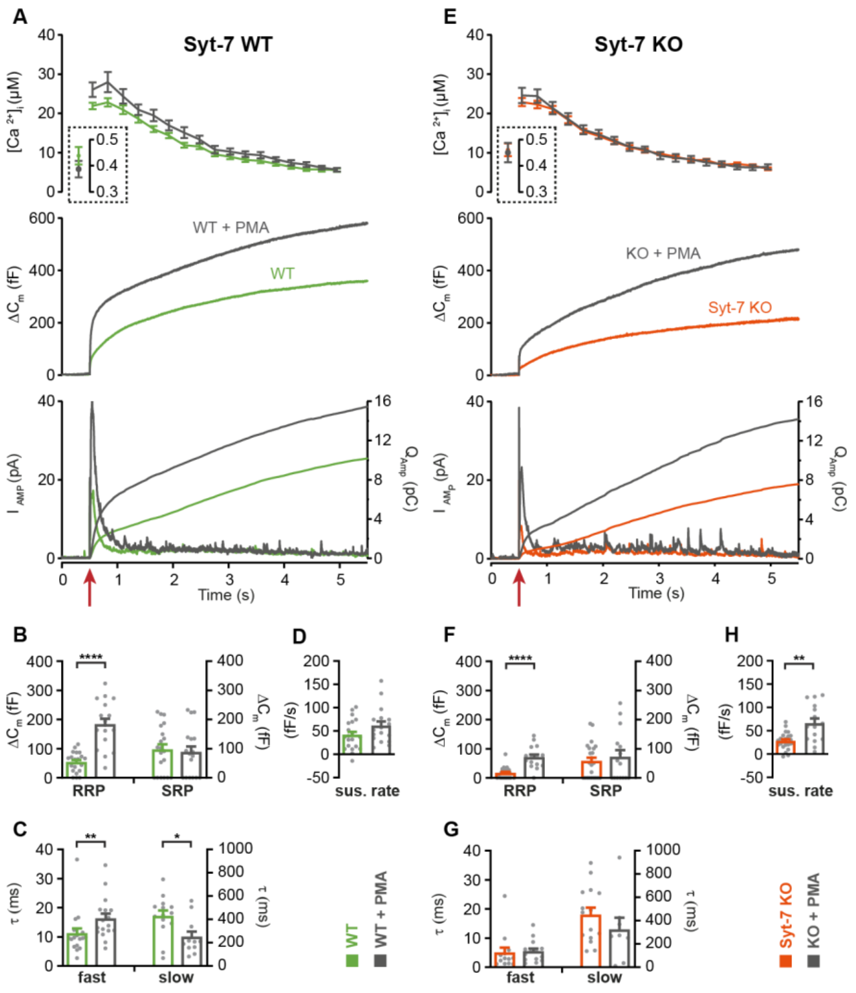


**Figure S5 – Application of phorbol esters to Syt-7 WT and KO at higher prestimulation [$Ca^{2+}$]**

**A**. Calcium uncaging experiment from high prestimulation [$Ca^{2+}$] in Syt-7 WT cells (green traces) and in Syt-7 WT cells perfused with 100 nM PMA (WT + PMA) (grey traces). Panels are arranged as in Fig. 1A. PMA treatment strongly augmented the primed pool size in WT cells.

**B.** Sizes of the RRP and SRP.

**C.** Time constants, $\tau$, of fusion for fast (i.e. RRP) and slow (i.e. SRP) secretion.

**D.** Sustained rates of secretion.

**E.** Calcium uncaging experiment from high prestimulation [$Ca^{2+}$] in Syt-7 KO cells (vermilion traces) and in Syt-7 KO cells perfused with 100 nM PMA (Syt-7 KO + PMA) (grey traces). PMA-induced potentiation of release was robust when Syt-7 KO cells were stimulated from higher prestimulation [$Ca^{2+}$].

**F.** Sizes of the RRP and SRP.

**G.** Time constants, $\tau$, of fusion for fast (i.e. RRP) and slow (i.e. SRP) secretion.

**H.** Sustained rates of secretion.

*: $p<0.05$; **: $p<0.01$; ****: $p<0.0001$, Mann-Whitney test. Number of cells: Syt-7 WT: N = 20 cells; Syt-7 WT + PMA: N = 17 cells; Syt-7 KO: N = 23 cells, Syt-7 KO + PMA: N = 15 cells.

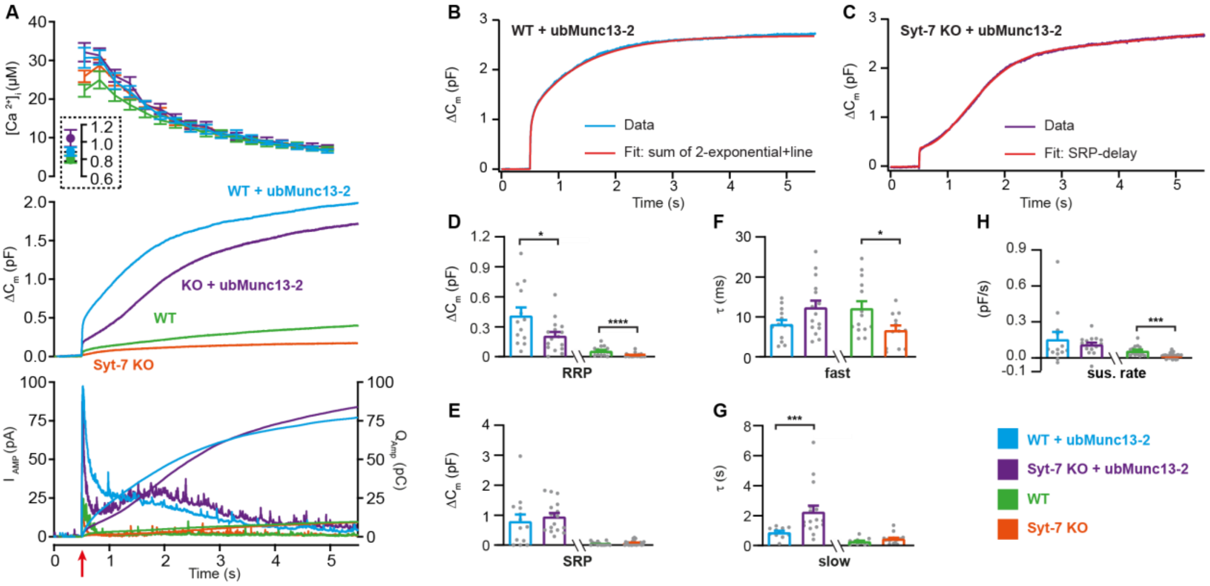


**Figure S6 – overexpressing ubMunc13-2 in Syt-7 WT and KO cells at higher prestimulation [$Ca^{2+}$].**

**A** Calcium uncaging experiment from high prestimulation [$Ca^{2+}$] in WT cells (green), WT overexpressing ubMunc13-2 (WT + ubMunc13-2) (cyan), Syt-7 KO (vermilion) and Syt-7 KO overexpressing ubMunc13-2 (KO + ubMunc13-2) (purple). Panels are arranged as in Fig. 1A. The overexpression ubMunc13-2 potentiated the release in both WT and Syt-7 KO cells with a remarkable delay of the SRP in the Syt-7 KO.

**B** A capacitance trace (‘Data’, blue) from a WT cell overexpressing ubMunc13-2 (cyan) fitted with a sum of two exponentials and a line (‘Fit’, red).

**C** A capacitance trace (‘Data’) from a Syt-7 KO cell overexpressing ubMunc13-2 (‘Fit’, purple) fitted with a modified function, taking into account the SRP-delay (red trace).

**D, E** Sizes of the RRP and SRP.

**F, G** Time constant, $\tau$, of fusion for fast (i.e. RRP) and slow (i.e. SRP) secretion. Note that the $\tau$ for the SRP in the Syt-7 KO + ubMunc13-2 group includes both the secretory delay and the fusion kinetics (Eq. 7) and is therefore not directly comparable to the $\tau$ from other groups.

**H** Sustained rate of secretion.

Data information: In (A-H) data with error bars are presented as mean ± SEM; in (A), the traces are the mean of all cells. WT (green) and Syt-7 KO (vermilion) are displayed here only to illustrate the massive increase upon ubMunc13-2 overexpression; statistical tests are only conducted for Syt-7 KO + ubMunc13-2 vs Syt-7 WT + ubMunc13-2 and Syt-7 WT vs Syt-7 KO. Statistics: *: $p<0.05$; **: $p<0.01$; ***: $p<0.001$; ****: $p<0.0001$. In D E, G and H): Mann-Whitney test. Number of cells: WT: N = 15 cells; WT + ubMunc13-2: N = 13 cells; Syt-7 KO: N = 25 cells; Syt-7 KO + ubMunc13-2: N = 16 cells.

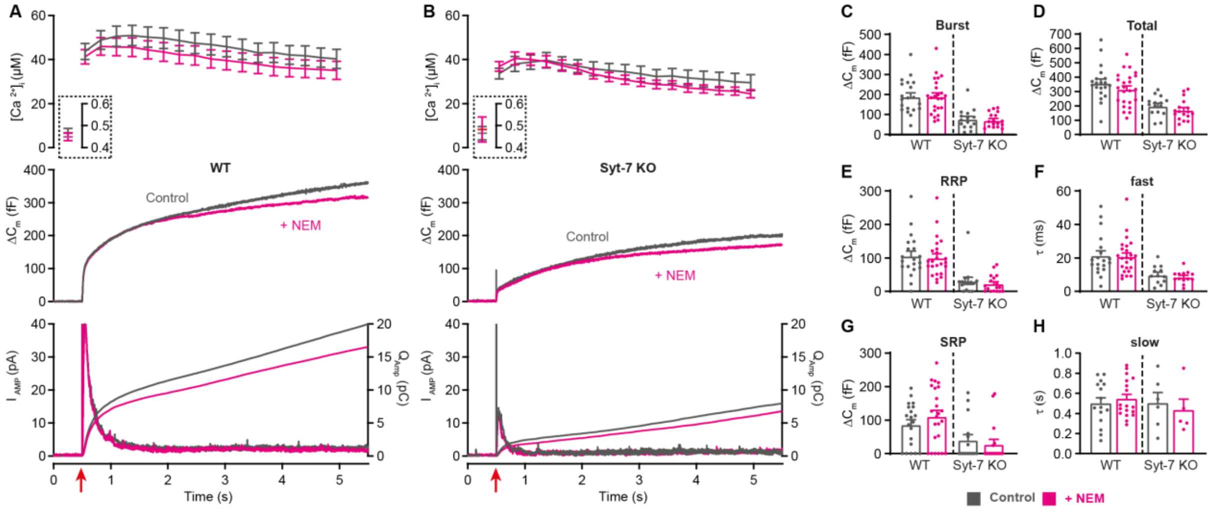


**Figure S7 – Blocking NSF had no effect in WT and Syt-7 KO cells when stimulated from high prestimulation [$Ca^{2+}$].**

**A,B** Calcium uncaging experiment from high prestimulation [$Ca^{2+}$] in WT (A) and in Syt-7 KO (B) control cells (Control, grey) and in cells infused with 200 µM N-Ethylmaleimide (+NEM, magenta). Panels are arranged as in Fig. 1A.

**C** Size of the burst (secretion within 0.5 s after the flash).

**D** Total release (secretion within 5 s after the flash).

**E** Size of the RRP.

**F** Time constant, $\tau$, of fusion for fast (i.e. RRP).

**G** Size of the SRP.

**H** Time constant, $\tau$, of fusion for slow (i.e. SRP).

Data information: Data are presented as mean ± SEM. *: $p<0.05$; **: $p<0.01$, Mann-Whitney test comparing control cells to cells infused with NEM from the same genotype. Number of cells: WT control: N = 20 cells; WT + NEM: N = 24 cells; Syt-7 KO control: N = 15 cells; Syt-7 KO + NEM: N = 17 cells.

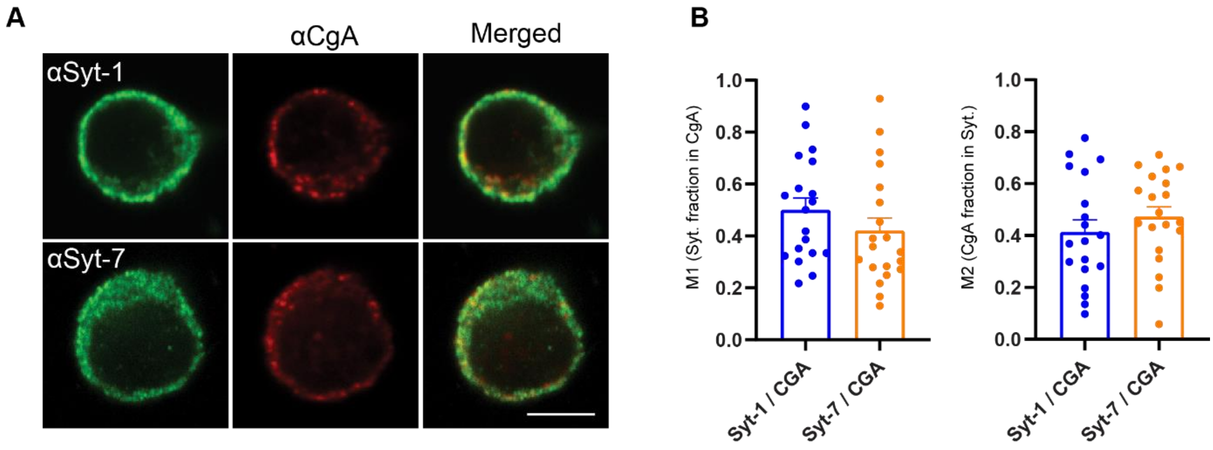


**Figure S8 – Synaptotagmin-1 and -7 co-staining with chromogranin-A (CgA).**

**A.** Single confocal slices of new-born mouse chromaffin cells stained against Syt-1 (α-Syt-1) or Syt-7 (α-Syt-7) and CgA (α-CgA) in WT cells, and merged images. Scale bar: 5 µm.

**B.** Manders' coefficients M1 and M2 (mean ± SEM) for the co-localization analysis of Syt-1 or Syt-7 and CgA in WT cells.

Data information: data with error bars are presented as mean ± SEM, Student t-test. Number of cells: Syt-1/CgA: N = 19 cells; Syt-7 CgA: N = 20 cells.

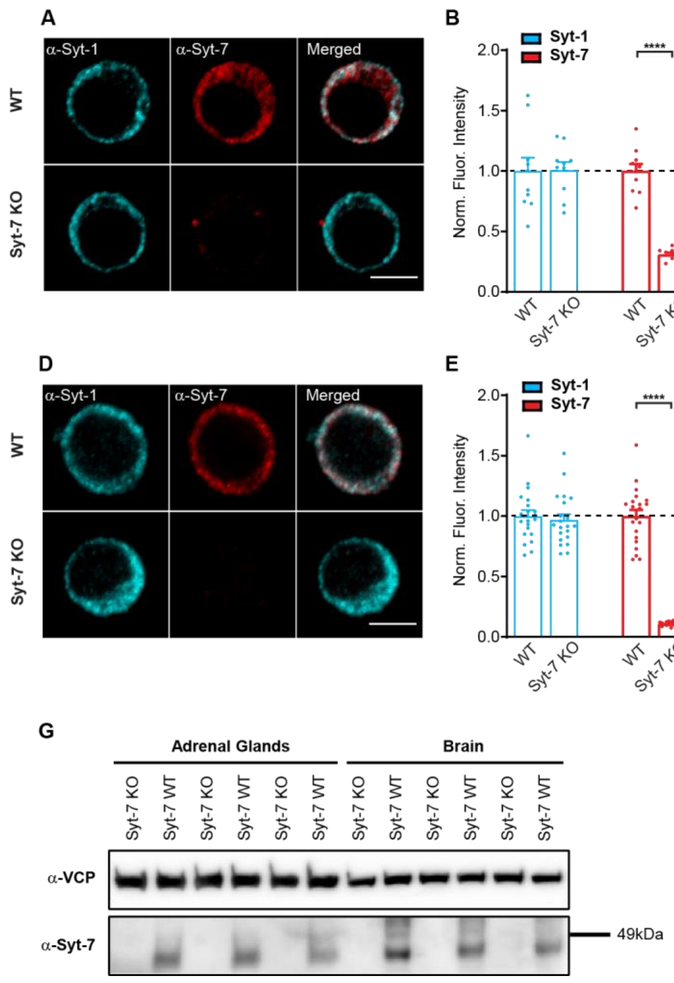


**Figure S9- Syt-1 and Syt-7 limited co-localization.**

A-C. Immunostaining and quantification of Syt-1 and Syt-7 expression in the absence of glutaraldehyde during the fixation step. Antibodies are a Syt-1 rabbit antibody and a Syt-7 mouse monoclonal antibody (both from Synaptic Systems; see Materials and Methods).

A. Single confocal slices of newborn mouse chromaffin cells stained against Syt-1 (α-Syt-1) and Syt-7 (α-Syt-7) in WT cells and in Syt-7 KO cells, and merged images. Scale bar: 5 µm.

B. Quantification of Syt-1 and Syt-7 staining in WT and Syt-7 KO cells. Some residual unspecific staining was detected in the Syt-7 KO cells.

C. Manders' coefficients (mean ± SEM) for the co-localization analysis of Syt-1 fraction in Syt-7 and Syt-7 fraction in Syt-1 in WT and Syt-7 KO cells.

D-F. Immunostaining and quantification of Syt-1 and Syt-7 expression using a mouse monoclonal Syt-7 antibody (clone 275/14, Sigma-Aldrich) and a rabbit polyclonal Syt-1 antibody (W855; T.C. Südhof).

D. Single confocal slices of newborn mouse chromaffin cells stained against Syt-1 ($\alpha$-Syt-1) and Syt-7 ($\alpha$-Syt-7) in WT cells and in Syt-7 KO cells, and merged images. Scale bar: 5 µm.

E. Quantification of the staining against Syt-1 and Syt-7 in WT and Syt-7 KO cells.

F. Manders' coefficients (mean ± SEM) for the co-localization analysis of Syt-1 fraction in Syt-7 and Syt-7 fraction in Syt-1 in WT and Syt-7 KO cells.

G. Protein immunoblot probed with antibodies against Syt-7 ($\alpha$-Syt-7) and VCP ($\alpha$-VCP) as a loading control in Syt-7 KO or WT adrenal glands and brains obtained from three new-born mice (one lane per animal). Syt-7 mouse monoclonal antibody (from Synaptic Systems; see Materials and Methods).

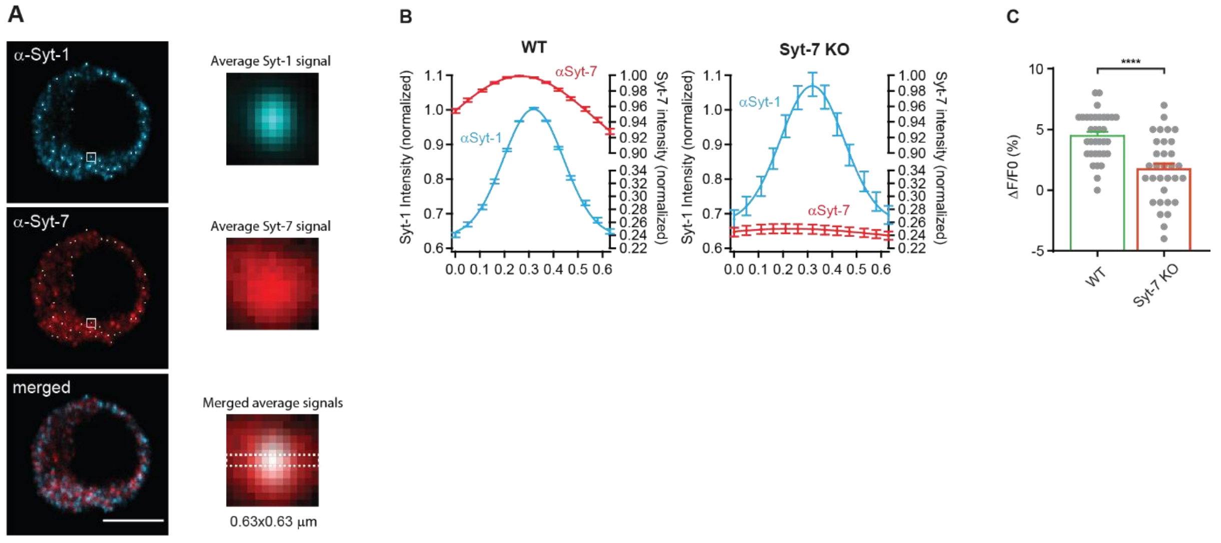


**Figure S10 – Averaging around Syt-1 peaks reveales an accumulation of Syt-7 signal colocalizing with Syt-1.**

**A** Left panel: An automatized averaging routine was used to identify peaks in the Syt-1-positive channel (white dots), followed by averaging in a square ROI surrounding each dot in both channels. Scale bar: 5 µm. Right panel: averaged vesicles from the Syt-1 and Syt-7 channel, and (bottom) after merging both channels.

**B** Line profiles (and Gaussian fits) through the equatorial plane of averaged images from WT (left panel) and Syt-7 KO (right panel) cells. The Syt-1 peak is sharper than the Syt-7 peak (see also average vesicles in panel A), because selection of peaks (vesicles) were made and aligned in the Syt-1 channel. Note the absence of a peak in the Syt-7 line profile in the Syt-7 KO.

**C** Syt-7 fluorescence change from 0.0 µm to 0.3 µm (peak) in WT and Syt-7 KO.

Data information: data with error bars are presented as mean ± SEM; Statistics: ****: $p<0.0001$, Mann-Whitney test. Number of cells in B-C: Syt-7 WT: 34 cells; Syt-7 KO: N = 32 cells.

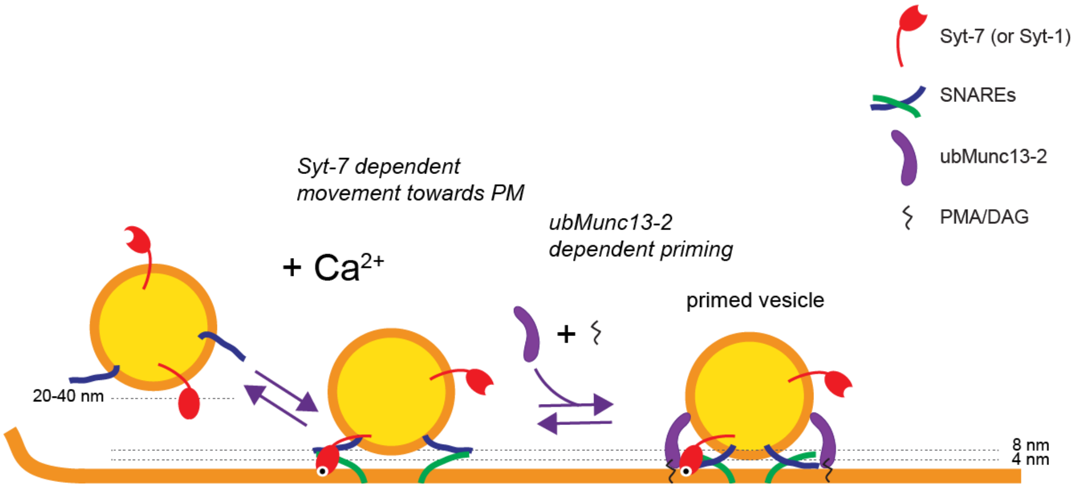


**Figure S11. Proposed role of Syt-7 and ubMunc13-2 in dense-core vesicle priming.**

Syt-7 reinforces recruitment of vesicles into a critical distance (6-10 nm) for ubMunc13-2 to bridge vesicle and plasma membrane for SNARE-complex formation. The ubMunc13-2 dependent priming-step is promoted by phorbol esters (PMA/DAG) and can be stabilized by blocking NSF-dependent de-priming. At high prestimulation $Ca^{2+}$, other molecular calcium-sensors likely contribute to $Ca^{2+}$-dependent recruitment of vesicles to the plasma membrane via a Syt-7 independent mechanism.
